# Two-Phase Coding Strategy by CA1 Pyramidal Neurons: Linking Spatiotemporal Integration to Predictive Behavior

**DOI:** 10.1101/2025.03.16.643530

**Authors:** Raphael Heldman, Dongyan Pang, Colin Porter, Xiaoliang Zhao, Alex Roxin, Yingxue Wang

**Affiliations:** Max Planck Florida Institute for Neuroscience, One Max Planck Way, Jupiter, FL 33458, USA; Graduate Program in Neuroscience, Boston University, 111 Cummington Mall, Boston, MA 02215, USA; Centre de Recerca Matemàtica, Campus de Bellaterra, Edifici C, 08193 Bellaterra (Barcelona), Spain

## Abstract

Space and time are fundamental components of memory, yet how the brain encodes these dimensions to guide behavior remains unclear. Using virtual-reality environments, we uncovered a two-phase neural code in hippocampus CA1 that represents time or distance through two functional pyramidal subpopulations, PyrUp and PyrDown. In Phase I, PyrUp activity synchronously increases to mark the initiation of encoding; In Phase II, their activity decays at heterogeneous, neuron-specific rates, creating a gradual divergence in across-population firing rates that scales with elapsed time. Conversely, PyrDown activity initially decreases before gradually rising. The crossover point, where rising PyrDown activity surpasses declining PyrUp activity, precedes predictive licking behavior. Combining optogenetics and computational modeling, we provided circuit-level evidence that PyrUp neurons primarily process locomotion-related inputs regulated by somatostatin-positive interneurons, whereas PyrDown neurons mainly receive reward-related inputs gated by parvalbumin-positive interneurons. These findings advance our understanding of how hippocampal circuits compute spatiotemporal information to inform behavior.

## INTRODUCTION

Our daily experiences, like finding a place to park the car near a restaurant at dinner time, are encoded into memories with embedded spatial and temporal contexts^1^. The hippocampus plays a central role in this encoding process^2–4^. Early research discovered “place cells” that activate when an animal enters specific locations^5^, forming neural maps that may provide spatial context for memories^6^. More recently, studies identified “time cells” that fire at successive moments during an experience^7–12^, potentially allowing chronological organization of events^13^. Both place and time cells maintain their specific tuning to distance or time even without external cues, exhibiting internally generated fields (IGFs). These IGFs activate sequentially, forming internally generated sequences (IGSs) to track traveled distance or elapsed time^7–12^.

However, only a subset of active pyramidal neurons exhibit precise temporal tuning, raising the question of whether the broader population of neurons can also encode spatiotemporal information. Recently, we uncovered two functional subpopulations of CA1 pyramidal neurons, PyrUp and PyrDown neurons, that exhibit two-phase dynamics potentially encoding elapsed time or traveled distance^14^. In phase I, PyrUp neurons synchronously increase firing rates, signaling the onset of encoding. In phase II, PyrUp activity decays at heterogeneous, neuron-specific rates, translating a gradual divergence of firing rates across the population into a metric of elapsed time or distance. In contrast, PyrDown neurons show an opposite pattern, where the activity shuts down in phase I and gradually ramps up in phase II. For both subpopulations, the two-phase dynamics potentially combine a “start” signal with subsequent time or distance encoding, extending beyond the classical sequential activation of place cells and IGF-expressing neurons. However, it remains unknown how hippocampal circuits generate these dynamics, and how these dynamics may inform behavior.

Local inhibitory interneurons are critical architects of hippocampal computation. Parvalbumin-positive (PV) and somatostatin-positive (SST) interneurons—two dominant classes in CA1— regulate pyramidal neuron activity through complementary pathways^15^. A major population of PV interneurons targets perisomatic regions, imposing fast and robust inhibition that synchronizes pyramidal cell output spikes^15–20^. In contrast, a major population of SST interneurons innervates apical dendrites, gating entorhinal cortical (EC) inputs while indirectly disinhibiting CA3 inputs^18,21–24^. This dual regulation positions SST interneurons to coordinate EC and CA3 input pathways during memory encoding. Our prior findings through optogenetic inactivation suggest that PV interneurons preferentially impact PyrDown dynamics while SST interneurons primarily regulate PyrUp dynamics^14^. However, how different excitatory input pathways influence these dynamics remains unclear.

In this study, we investigated two interconnected questions: (1) How do PyrUp and PyrDown dynamics link to behavior? and (2) How do hippocampal circuits shape these dynamics? Combining cell-type-specific activation of inhibitory interneurons with computational modeling, we hypothesized a simplified circuit model: PyrUp neurons mainly receive locomotion-driven dendritic inputs regulated by SST interneurons, while PyrDown neurons primarily process reward-related perisomatic inputs gated by PV interneurons. PyrUp and PyrDown activity together informs the prediction of reward. Collectively, our findings offer new insights into how hippocampal circuits transform abstract neuronal dynamics into behavior.

## RESULTS

### CA1 pyramidal neurons exhibit ramping activity during distance or time integration

We trained mice in a virtual reality-based path integration (PI) task that required mice to estimate traveled distance to obtain rewards, as previously described^14^ (Fig. 1a). In each trial, a brief sensory cue appeared on the screens, after which the mouse was required to run 180 cm while the screens displayed a uniform gray background, creating a cue-constant segment. The animal then had to actively lick within a designated reward zone to trigger a drop of water reward; failure to do so resulted in no reward (Fig. 1b). After training, animals developed consistent behavioral patterns, beginning to accumulate distance by initiating running independently of the visual cue that marked trial start^14^ (Fig. 1c). Their running speed profiles were highly consistent across trials (Fig. 1c), making it difficult to distinguish whether animals integrated over distance or time.

**Figure 1.**
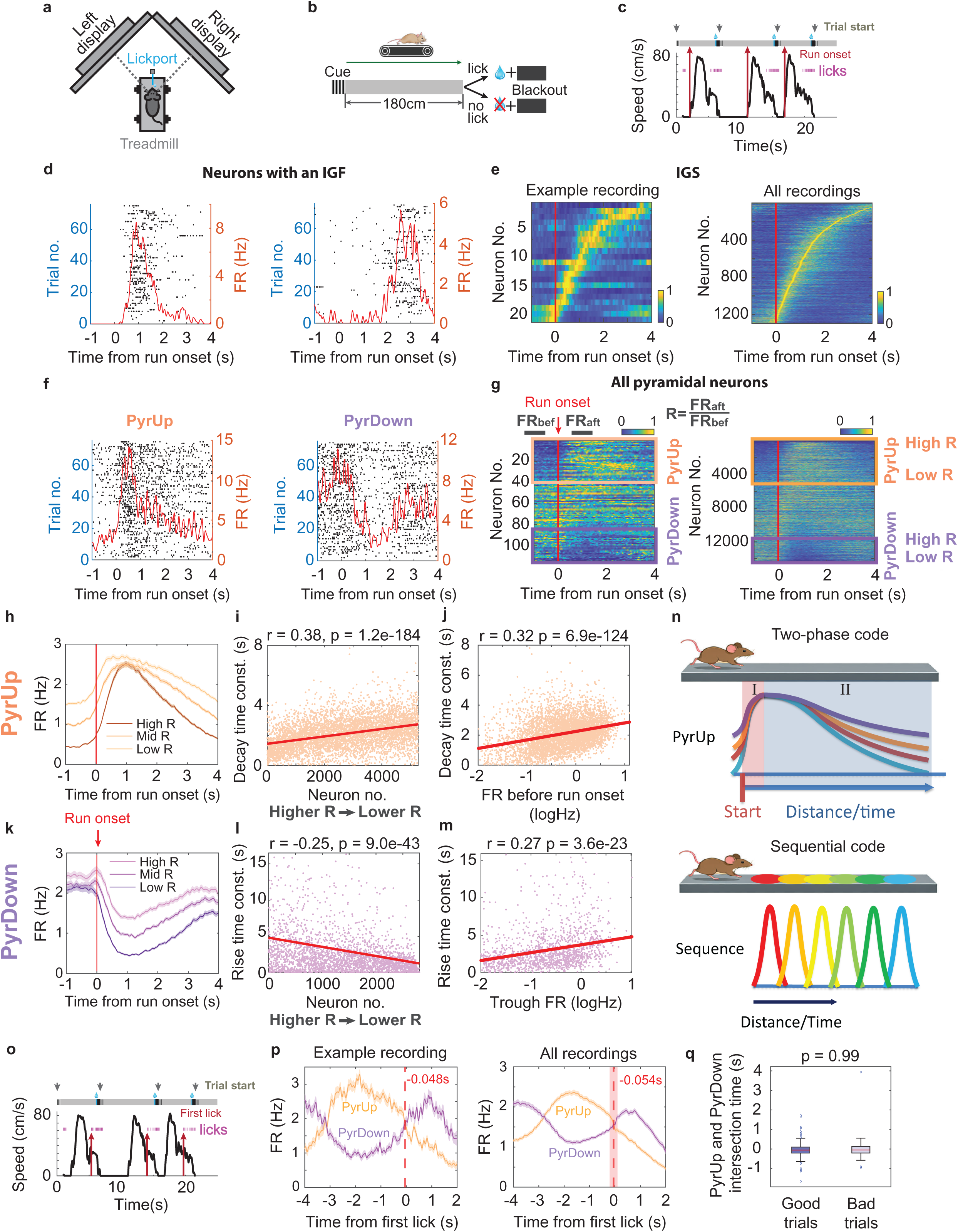
CA1 pyramidal neuron dynamics during a path integration task. (a) Virtual reality setup: Schematic of a head-fixed mouse with eye alignment to two monitors. The mouse runs on a treadmill and receives water reward through a lick port. (b) Task schematics for the PI task (Methods). (c) Speed (black) and licking (magenta) over time across three consecutive trials. Run onsets (red arrows) and cue onsets (gray arrows) are temporally dissociated. (d) Spike rasters and average firing rate profile (red line) for two IGF-expressing pyramidal neurons with activity aligned to run onset. Data from a single recording session during the PI task. (e) Left: Heatmap of an IGS from one recording. Right: Aggregated IGS across all IGF-expressing neurons across all recordings. (f) Spike rasters and averaged firing rate profiles (red line) for example PyrUp (left) and PyrDown (right) neurons from the same session. (g) Left: Heatmaps of all pyramidal neurons (one recording) ordered by ratio R = FR_aft_/FR_bef_ (FR_bef_: baseline rate [−1.5 to −0.5s]; FR_aft_: post-run rate [0.5 to 1.5s]; 0s = run onset). Right: Neurons pooled across recordings (≥15 good trials), with PyrUp (5,273; orange) and PyrDown (2,808; purple) subgroups highlighted. (h) Firing rate profiles (±SEM) for PyrUp neurons stratified into high/mid/low R tertiles (dark to light orange). (i) Post-peak decay time constants (orange dots) for PyrUp neurons (ordered by descending R). Red line: linear regression (see Methods). (j) The baseline firing rate before run onset (log(FR_bef_)) versus decay time constant for PyrUp neurons. Red line: linear regression. (k-m) Analogous to panels (h–j) for PyrDown neurons (purple). (n) Cartoon illustrating the two-phase code by PyrUp neurons (top) versus the sequential code by place cells or IGF-expressing neurons (bottom). (o) Speed and licking traces (as in panel (c)) with first predictive licks per trial highlighted (red arrows). (p) Left: first-predictive-lick aligned firing rate profile averaged across all PyrUp (orange) and PyrDown (purple) neurons from one example recording. Red dotted line: PyrUp and PyrDown activity intersection. Right: Aggregated data from all the recordings. Median intersection time (red dotted line) ± MAD (shading). (q) Intersection time distributions in good versus bad trials.

Using silicon probes, we recorded from the hippocampal CA1 region. When aligning neuronal activity to run onset, we identified a subset of pyramidal neurons that exhibited firing fields at specific time points after run onset. Given that the visual cues remained constant, these firing fields represented internally generated fields (IGFs) (Fig. 1d, mean ± SEM = 9.05 ± 0.50%, 1295 of 14306 neurons, n = 53 animals, 192 recordings. Mean ± SEM is used if not specifically mentioned). The IGF-expressing neurons activated sequentially to form an internally generated sequence (IGS) that spanned the integration period (Fig. 1e), consistent with our previous observations^14^.

Since IGF-expressing neurons constituted a small fraction of the pyramidal population, we sought to determine whether the broader neuronal population encoded spatial or temporal information during the integration period. Analysis of the entire pyramidal population (14306 neurons) uncovered two distinct functional subpopulations with complementary responses around run onset and elapsed time that can be decoded using their dynamics^14^. These subpopulations together comprised over half of the active CA1 pyramidal neurons. One group, termed “PyrUp” neurons (36.86% of pyramidal neurons), rapidly increased their activity at run onset and exhibited gradual decay as the animal approached the reward zone (Fig. 1f left and 1g). The other group, termed “PyrDown” neurons (19.63% of pyramidal neurons), initially shut down their activity before gradually ramping up toward the reward (Fig. 1f right and 1g). Unlike the precisely tuned IGF- expressing neurons, most PyrUp and PyrDown neurons exhibited gradual changes in firing rate without precise temporal tuning^14^ (Fig. 1f,g).

### A Two-Phase Coding Mechanism for time or distance

Our previous work proposed that PyrUp and PyrDown neurons implement a two-phase neural code to represent elapsed time or distance, instead of merely tracking running speed^14^. For PyrUp neurons, in phase I, they exhibited a synchronous increase in their activity at run onset, reaching their peak activity at similar times across the population (Fig. 1h, peak time: 1.19 ± 0.01 s). This synchronized increase marks the initiation of encoding. In phase II, PyrUp neurons decayed from their peak firing rates at heterogeneous, neuron-specific rates. This heterogeneity in decay rates creates a gradual divergence in firing rates across the population, with greater divergence corresponding to longer elapsed times (Fig. 1h).

Neurons with slower decay rates tended to exhibit higher baseline firing before run onset and smaller increases in activity around run onset (lower ratio R=FR_aft_/FR_bef_, where FR is firing rate, Fig. 1i,j). This relationship was observed across all recorded pyramidal neurons (Fig. 1i) and within individual recordings, with 55.9% of recording sessions showing significant correlations between the increase in activity around run onset and the decay rate.

PyrDown neurons displayed dynamics opposite to those of PyrUp neurons. Their activity transiently shut down at run onset, reaching a trough at similar time points (1.15 ± 0.01s). Following the trough, PyrDown activity gradually ramped up at neuron-specific rates. Faster rises correlated with lower baseline activity measured at the trough and larger decreases in activity around run onset (lower R values) (Fig. 1k-m).

Overall, the heterogeneity in PyrUp decay and PyrDown rise rates enables a population-level coding strategy for the passage of time (Fig. 1n). Unlike classical sequential coding by place cells or IGF-expressing neurons, which are precise tuned to specific distance or time points, this strategy encodes elapsed time through gradual, monotonic changes in firing rates across the population, without requiring precise tuning in individual neurons (Fig. 1n).

### Behavioral relevance of PyrUp and PyrDown dynamics

We then investigated how the PyrUp and PyrDown dynamics relate to animals’ behavior in the PI task. These subpopulations displayed distinct behavioral associations: PyrUp neurons exhibited higher activity during running, while PyrDown neurons became more active as animals approached the reward zone.

Expanding beyond two-phase dynamics, we further analyzed how PyrUp and PyrDown activity aligns with key task-related behavioral events. We identified neurons significantly tuned to each of three behavioral events—run onset, the first predictive lick, and reward delivery—by comparing their activity to shuffled data (Supplementary Fig. 1a). PyrUp neurons were predominantly associated with the run-related event, with 22% showing significant tuning to run onset. In contrast, only 5% of PyrDown neurons exhibited such tuning. Conversely, PyrDown neurons were preferentially linked to reward-related events: 15% were tuned to reward delivery and 8% to the first predictive lick. In contrast, only 1% and 3% of PyrUp neurons, respectively, were tuned to these events (Supplementary Fig. 1b).

These results suggest a functional division: PyrUp neurons primarily process run-related inputs, engaged during running and time or distance tracking; whereas PyrDown neurons tend to receive reward-related inputs, recruited when rewards are anticipated and consumed.

### Intersection time of PyrUp and PyrDown dynamics signals imminent reward

Our findings suggest that PyrUp and PyrDown subpopulations implement a two-phase code for tracking elapsed time and exhibit different behavioral associations. To investigate how their dynamics inform behavior, such as predicting the upcoming reward, we analyzed their relationship to predictive licking—a behavior marker of reward anticipation. Aligning neuronal activity to the first predictive lick revealed that the intersection time between the rising PyrDown activity and declining PyrUp activity in phase II preceded the licking onset (Fig. 1p left). This temporal relationship was robust across recordings (Fig. 1p right, Median ± MAD = −54 ± 153ms). This relationship persisted when using an alternative method to identify PyrUp and PyrDown neurons (validated via shuffled activity around run onset; Supplementary Fig. 2e, Median ± MAD = −119 ± 162ms, Methods).

Next, we determined whether the neural-behavioral relationship holds regardless of whether animals accurately estimated time or distance. We separate trials into two categories. Trials with accurate integration (good trials) met the following criteria: a complete stop before run onset, largely uninterrupted running, and successful reward acquisition. These trials constituted 82.1±1.7% of all trials (100% reward rate). In contrast, in trials that do not fulfill all these criteria (bad trials) (82.8±4.2% reward rate), animals initiated licking significantly earlier (Supplementary Fig. 2a) despite comparable mean running speed (Supplementary Fig. 2b). In bad trials, both PyrUp and PyrDown subpopulations exhibited attenuated firing rate changes around run onset (Supplementary Fig. 2c-d)^14^. However, the temporal alignment between the intersection of PyrUp/PyrDown activity and the first predictive lick remained unchanged in both trial types (Fig. 1q and Supplementary Fig. 2f).

These results showed that the intersection where PyrDown activity surpasses PyrUp may serve as a signal for imminent reward, playing a role in informing predictive licking even when animals misjudge time or distance.

### Inactivating pyramidal neuron activity at run onset does not affect integration accuracy

To determine whether PyrUp and PyrDown dynamics causally contribute to the accuracy of distance or time integration, we selectively suppressed pyramidal neuron activity during each phase of the two-phase code. To transiently silence CA1 pyramidal neurons, we optogenetically activated PV interneurons, which primarily exert rapid perisomatic inhibition over pyramidal neurons^25^.

First, we targeted phase I, the encoding initiation phase. In PV-Cre mice, we transfected dorsal CA1 with channelrhodopsin-2 (ChR2)^26^ and recorded unilateral CA1 neuronal activity using an optrode. Delivering a 0.5-second light pulse at run onset (Fig. 2a, Supplementary Fig. 5b-c) transiently suppressed firing rates in both PyrUp and PyrDown neurons (Fig. 2b-2e). Activity recovered quickly after stimulation (mean FR 1 to 1.5 s: PyrUp: control: 2.56 ± 0.13Hz, stimulation: 2.39 ± 0.13Hz, p = 0.07; PyrDown: control: 1.18±0.10Hz, stimulation: 1.10±0.09Hz, p = 0.48, Wilcoxon rank-sum test is used if not specifically mentioned). Despite the suppression, phase II dynamics—characterized by heterogenous PyrUp decay and PyrDown rise rates—remained largely intact (PyrUp decay rate: control: 2.31 ± 0.05s, stimulation: 2.25 ± 0.06s, p = 0.45; PyrDown rise rates: control: 3.95 ± 0.32s, stimulation: 3.23 ± 0.33s, p = 0.12, two-sample t-test). IGF-expressing neurons also exhibited a transient firing rate decrease and reduced across-trial activity correlation (Fig. 2f-g).

**Figure 2.**
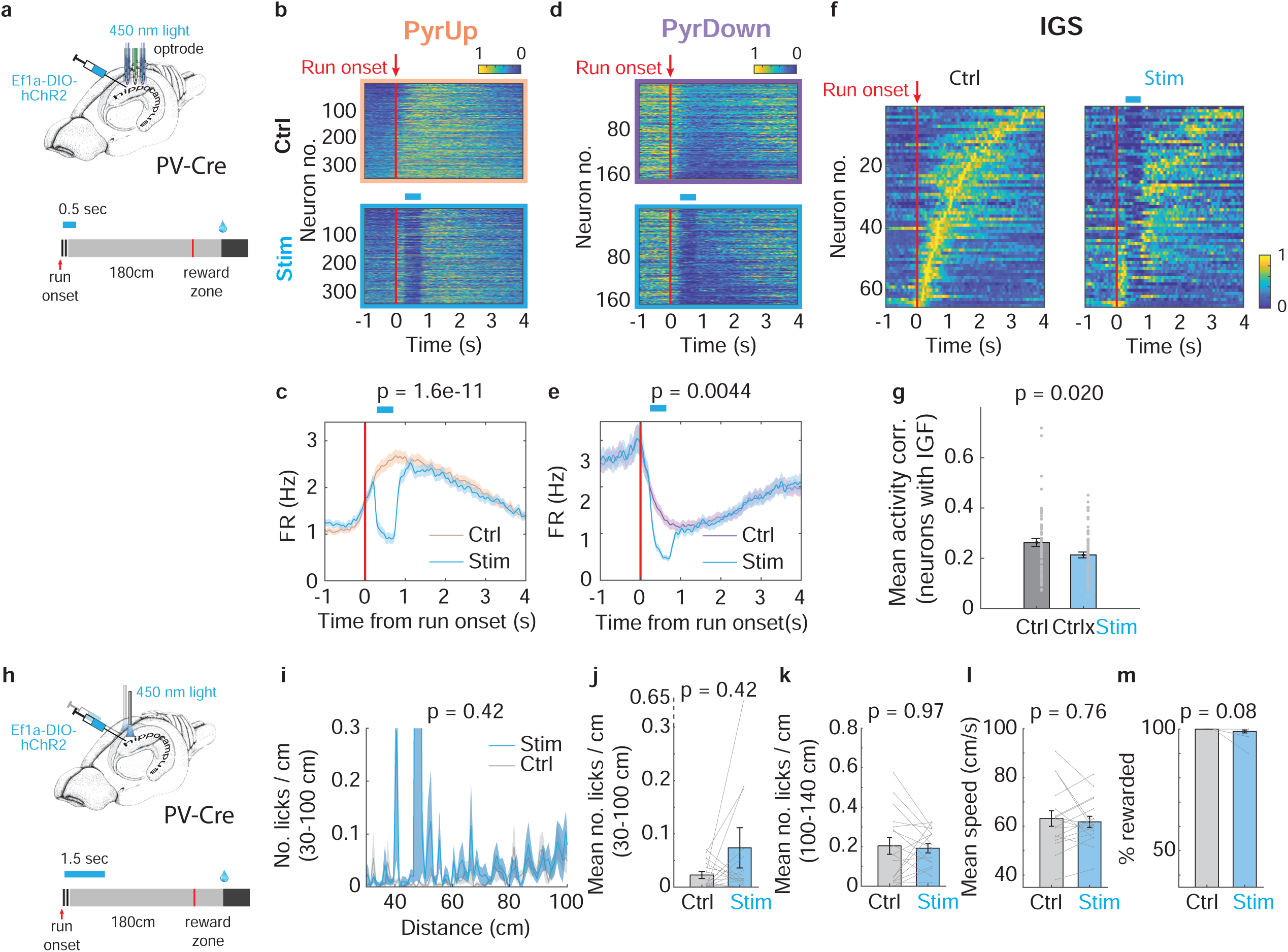
Pyramidal neuron inactivation at run onset. (a) PV interneuron activation protocol: Optogenetic stimulation (0.5 s duration, starting at run onset) delivered via optrode in a subset of trials (see Methods). (b) Heatmaps of PyrUp neurons during control (top) and stimulation trials (bottom). Data pooled from 5 animals across 13 recordings (342 of 928 neurons). Neurons ordered by firing rate ratio (R = FRaft/FRbef). Blue bar: estimated stimulation window. (c) PyrUp average firing rate profiles for control (orange) and stimulation trials (blue). Wilcoxon rank-sum test (FR 0–1s): control = 2.39 ± 0.12 Hz, stimulation = 1.49 ± 0.10 Hz, p = 1.6e-11. (d) Analogous to panel (b), for PyrDown neurons (166 of 928 neurons). (e) Analogous to panel (c), for PyrDown neurons. FR 0-1s: control: 1.63 ± 0.14Hz, stimulation: 1.19 ± 0.11Hz, p = 0.0044. (f) IGS during control (left) and stimulation trials (right). (g) Mean activity correlation among IGF-expressing neurons. Control = 0.26 ± 0.02, stimulation = 0.21 ± 0.01, p = 0.02. (h) Experimental setup to assess the behavioral effects of PV activation. The same protocol as panel (a), in a separate cohort of animals with bilateral optic fiber implants. (i) Lick histogram from the first 100 cm for control (grey) and stimulation trials (blue). Data from 6 animals, 17 recordings. (j-m) (j) Mean number of licks/cm from the first 100 cm based on panel (i); (k) Mean number of licks/cm between 100-140 cm; (l) Mean running speed; (m) Percentage of rewarded trials for control and stimulation trials.

To evaluate the behavioral impact of this manipulation, we performed bilateral optogenetic activation of PV interneurons during the PI task in a separate cohort of animals (Fig. 2h). PV activation at run onset did not significantly affect the licking pattern, mean running speed, or the percentage of rewarded trials. Task performance remained largely unchanged even when extending the stimulation duration from 0.5 s to 1.5 s (Fig. 2i-m).

Overall, these results suggest that the PyrUp and PyrDown activity during phase I, where the neurons synchronously increase (PyrUp) or decrease (PyrDown) activity, is not necessary for the integration accuracy.

### Inactivating pyramidal neuron activity mid-trial impairs integration accuracy

To test whether phase II, the time-encoding phase, of PyrUp and PyrDown dynamics is responsible for integration accuracy, we optogenetically activated PV interneurons starting at 120 cm into the cue constant segment for 1.5s (Fig. 3a). This manipulation significantly suppressed the ramping activity of PyrUp neurons (Fig. 3b-c) and PyrDown neurons (Fig. 3d-e). PyrUp decay rates were significantly reduced (control: 1.89 ± 0.05s, stimulation: 1.43 ± 0.06s, p = 3.24e-09, two-sample t-test), while PyrDown rise rates remained largely intact (control: 4.47 ± 0.94s, stimulation: 4.03 ± 0.93s, p = 0.74, two-sample t-test). IGF-expressing neurons showed no significant change to this manipulation (Fig. 3f-g).

**Figure 3.**
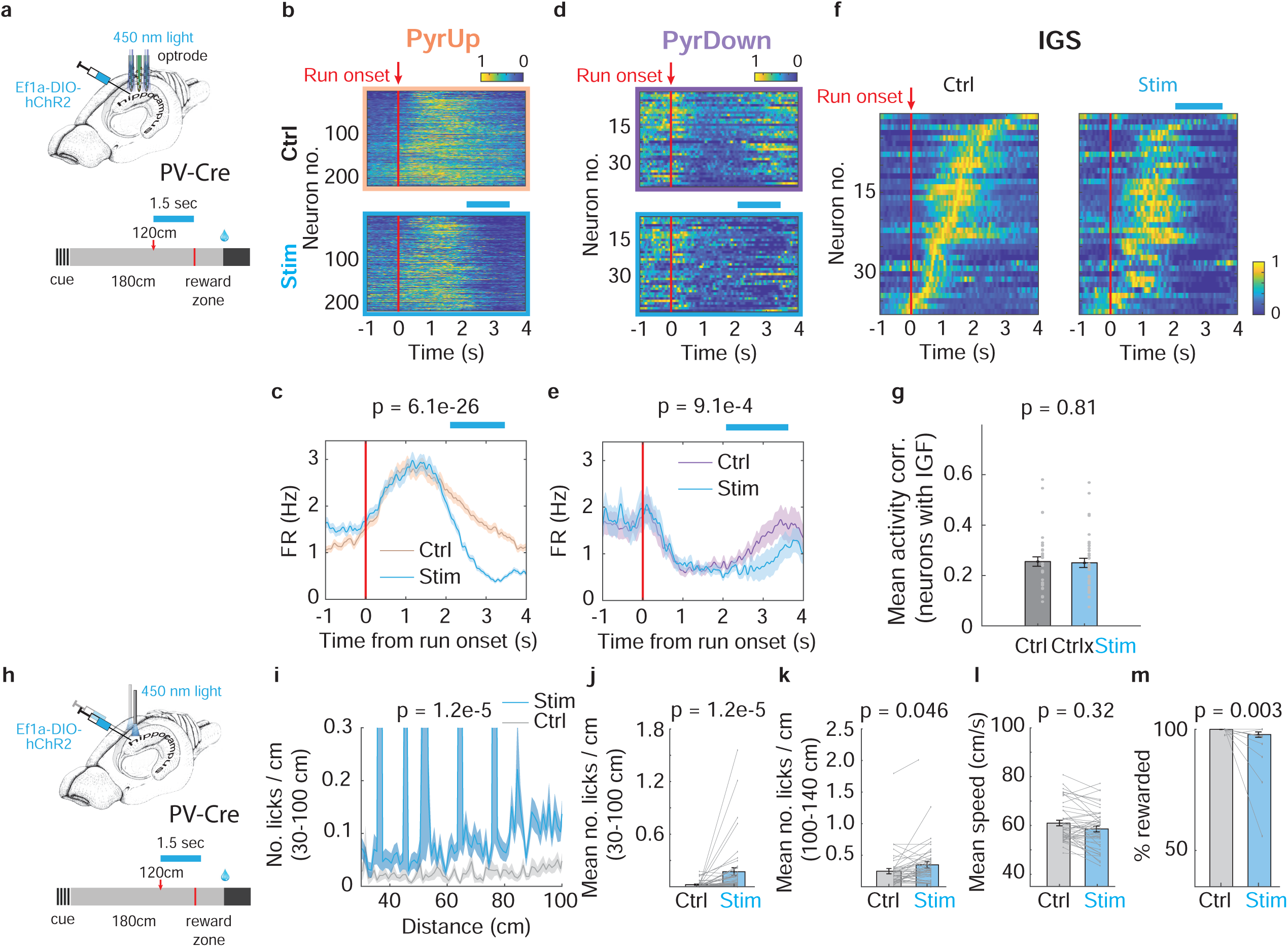
Pyramidal neuron inactivation in the middle of a trial. (a) PV interneuron activation protocol: Optogenetic stimulation (1.5 s duration, starting at 120 cm during the cue-constant segment) delivered via optrode in a subset of trials (see Methods). (b) Heatmaps of PyrUp neurons during control (top) and stimulation trials (bottom). Data pooled from 2 animals across 6 recordings (216 of 441 neurons). Neurons ordered by firing rate ratio (R = FRaft/FRbef). Blue bar: estimated stimulation window. (c) PyrUp average firing rate profiles for control (orange) and stimulation trials (blue). Wilcoxon rank-sum test (FR 2.5-3.5s): control: 1.60 ± 0.09Hz, stimulation: 0.60 ± 0.05Hz, p = 6.1e-26. (d) Analogous to panel (b), for PyrDown neurons (39 of 441 neurons). (e) Analogous to panel (c), for PyrDown neurons. FR 2.5-3.5s: control: 1.34 ± 0.22Hz, stimulation: 0.79 ± 0.25Hz, p = 9.1e-4. (f) IGS during control (left) and stimulation trials (right). (g) Mean activity correlation among IGF-expressing neurons. Control = 0.26 ± 0.02, stimulation = 0.25 ± 0.02, p = 0.81. (h) Experimental setup to assess the behavioral effects of PV activation. The same protocol as panel (a), in a separate cohort of with bilateral optic fiber implants. (i) Lick histogram from the first 100 cm for control (grey) and stimulation trials (blue). Data from 6 animals, 46 recordings. (j-m) (j) Mean number of licks/cm from the first 100 cm based on panel (i); (k) Mean number of licks/cm between 100-140 cm; (l) Mean running speed; (m) Percentage of rewarded trials for control and stimulation trials.

To examine the behavioral consequence, we bilaterally activated PV interneurons in the same cohort of animals as in Fig. 2 at 120 cm in a subset of trials (Fig. 3h). This manipulation led to a significant decrease in the percentage of rewarded trials (Fig. 3m). The behavioral deficits persisted into the control trial immediately following stimulation, where animals exhibited a significant increase in early licking before 100 cm (Fig. 3i-j) and predictive licking between 100- 140 cm (Fig. 3k). However, the mean running speed did not significantly change during the stimulation trials (Fig. 3l) or in the subsequent control trials, ruling out locomotor confounds (Supplementary Fig. 3a-b). Control experiments further ruled out that the effects were due to laser light alone (Supplementary Fig. 4a-f).

Together, these results suggest that inactivating PyrUp and PyrDown activity during phase II impairs integration accuracy, supporting their necessity for behavioral performance.

### Activating SST interneurons at the run onset preferentially suppresses PyrUp neurons without affecting integration accuracy

While pyramidal neuron output spike suppression via PV activation revealed the necessity of distinct coding phases for task performance, this approach cannot delineate how specific excitatory afferent pathways shape PyrUp and PyrDown dynamics. In contrast, CA1 SST interneurons primarily innervate apical dendritic compartments, modulating inputs from the entorhinal cortex (EC) and hippocampal CA3 region^18,21–24^. A major subset of SST interneurons directly inhibit distal dendrites receiving EC inputs, and may also indirectly disinhibit proximal dendrites targeted by CA3 inputs by silencing intermediate radiatum interneurons^21,23^. Our previous work demonstrated that SST interneuron inactivation after run onset selectively attenuated PyrUp activity^14^. This finding supports the idea that SST inactivation indirectly suppresses CA3 inputs and thus diminishes PyrUp dynamics. However, the role of EC inputs in shaping PyrUp and PyrDown dynamics remains unclear.

To address this, we transfected CA1 SST interneurons with ChR2 in SST-IRES-Cre animals and optogenetically activated them unilaterally at run onset for 0.5s to target phase I (Fig. 4a, Supplementary Fig. 5d-e). SST activation significantly suppressed PyrUp activity (Fig. 4b-c). This suppression likely reflected inhibition of EC inputs, as most SST interneurons in these mice reside in the striatum oriens and project extensively to the striatum oriens-lacunosum moleculare (OLM) layer (Supplementary Fig. 5a). Nevertheless, we cannot rule out some contribution from few SST interneurons targeting proximal dendrites and thus suppressing CA3 inputs. (Supplementary Fig. 5a).

**Figure 4.**
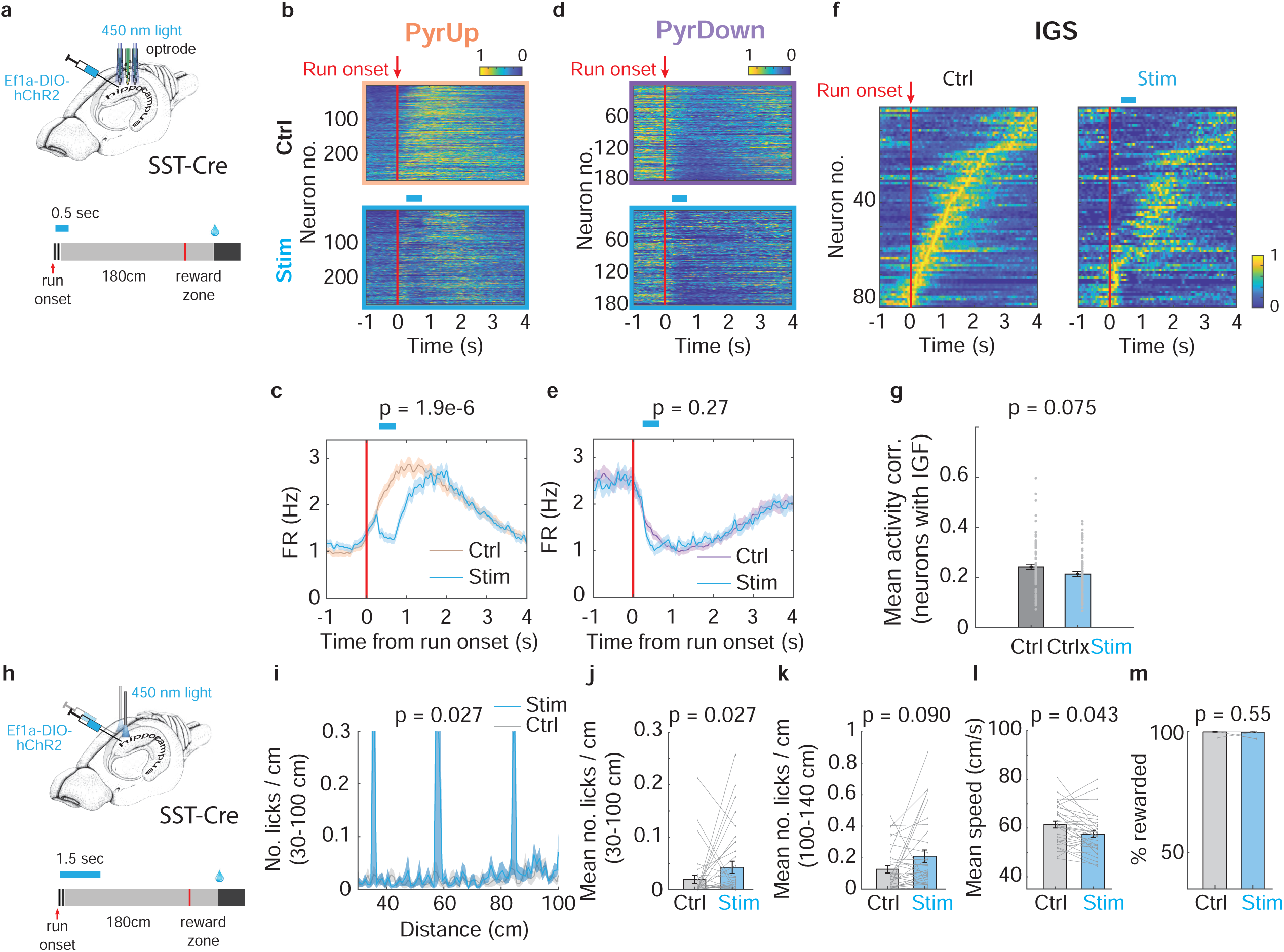
SST interneuron activation at run onset. (a) SST interneuron activation protocol: Optogenetic stimulation (0.5 s duration, starting at run onset) delivered via optrode in a subset of trials (see Methods). (b) Heatmaps of PyrUp neurons during control (top) and stimulation trials (bottom). Data pooled from 6 animals across 14 recordings (273 of 784 neurons). Neurons ordered by firing rate ratio (R = FRaft/FRbef). Blue bar: estimated stimulation window. (c) PyrUp average firing rate profiles for control (orange) and stimulation trials (blue). Wilcoxon rank-sum test (FR 0–1s): control: 2.25 ± 0.13Hz, stimulation: 1.51 ± 0.10Hz, p = 1.9e-6. (d) Analogous to panel (b), for PyrDown neurons (182 of 784 neurons). (e) Analogous to panel (c), for PyrDown neurons. FR 0-1s: control: 1.52 ± 0.11Hz, stimulation: 1.44 ± 0.11Hz, p = 0.27. (f) IGS during control (left) and stimulation trials (right). (g) Mean activity correlation among IGF-expressing neurons. Control = 0.24 ± 0.01, stimulation = 0.21 ± 0.01, p = 0.075. (h) Experimental setup to assess the behavioral effects of SST activation. The same protocol as panel (a), in a separate cohort of with bilateral optic fiber implants. (i) Lick histogram from the first 100 cm for control (grey) and stimulation trials (blue). Data from 5 animals, 33 recordings. (j-m) (j) Mean number of licks/cm from the first 100 cm based on panel (i); (k) Mean number of licks/cm between 100-140 cm; (l) Mean running speed; (m) Percentage of rewarded trials for control and stimulation trials.

The PyrUp suppression induced by SST activation was weaker than with PV activation which directly suppressed spikes at the soma (Firing rate change from control between 0 and 1 s: SST activation: −26.1 ± 3.4%, PV activation: −37.0 ± 2.4%, p = 0.0065), implying remaining afferent inputs sustaining PyrUp activity. Consistent with PV activation, PyrUp decay rates during phase II were not significantly altered (control: 2.21 ± 0.05s, stimulation: 2.15 ± 0.08s, p = 0.51).

In contrast, the PyrDown activity and their rise rates remained largely unaffected by SST activation (Fig.4d-e, rise rates: control: 3.39 ± 0.29s, stimulation: 3.26 ± 0.34s, p = 0.79, two-sample t-test). Cross-correlogram analysis^27^ did not reveal significant differences in monosynaptic connectivity from SST interneurons to PyrUp versus PyrDown neurons (SST to PyrUp: 0.012 ± 0.004; SST to PyrDown: 0.007 ± 0.003, p = 0.20), suggesting that preferential connectivity alone does not account for the selective suppression of PyrUp neurons. Instead, this differential effect may reflect differences in dendritic inputs, with the PyrDown neurons receiving weak excitation, particularly from EC, after run onset. Our previous findings showed that inactivating SST interneurons after run onset did not affect the PyrDown activity, suggesting their low CA3 inputs. Together, these data support that PyrDown neurons likely receive limited excitation from both EC and CA3 inputs during this period. Moreover, IGF-expressing neurons showed no significant change in trial-by-trial activity correlations during SST activation (Fig. 4f-g).

We further assessed the behavioral impact of SST interneuron activation by performing bilateral optogenetics at run onset for 1.5s in a separate cohort of animals (Fig. 4h). This manipulation produced only mild changes in early licking and running speed (Fig. 4i-l), with no significant effect on the percentage of rewarded trials (Fig. 4m, Supplementary Fig. 4g-l). These results are consistent with our PV activation, suggesting that neuronal activity in phase I is not critical for overall integration accuracy. The mild behavioral effects may result from the gradual recovery of PyrUp activity after stimulation (mean FR 1 to 1.5 s: PyrUp: control: 2.79 ± 0.15Hz, stimulation: 2.35 ± 0.15Hz, p = 0.003), which may overlap with the subsequent time-encoding phase II.

In summary, these SST activation results suggest that dendritic inputs, particularly the EC inputs, preferentially drive PyrUp activity during phase I. However, these inputs appear to have only a marginal impact on overall integration accuracy.

### Activating SST interneurons mid-trial preferentially suppresses PyrUp neurons without affecting integration accuracy

To investigate how SST interneurons regulate PyrUp and PyrDown activity during phase II, we optogenetically activated SST interneurons starting at 120 cm into the cue-constant segment for 1.5s (Fig. 5a). Similar to run-onset stimulation, this manipulation significantly suppressed PyrUp activity (Fig. 5b-c), again to a lesser extent than PV activation (firing rate change from control between 2.5 and 3.5 s, (%): SST activation: −41.5 ± 2.9%, PV activation: −60.0 ± 2.7%, p = 4.2e-9). These results further support the role of dendritic inputs, particularly EC inputs, in maintaining PyrUp activity. In addition, while PV activation markedly reduced PyrUp decay rates, SST activation caused a mild reduction (control: 2.14 ± 0.06s, stimulation: 1.90 ± 0.09s, p = 0.03, two-sample t-test).

**Figure 5.**
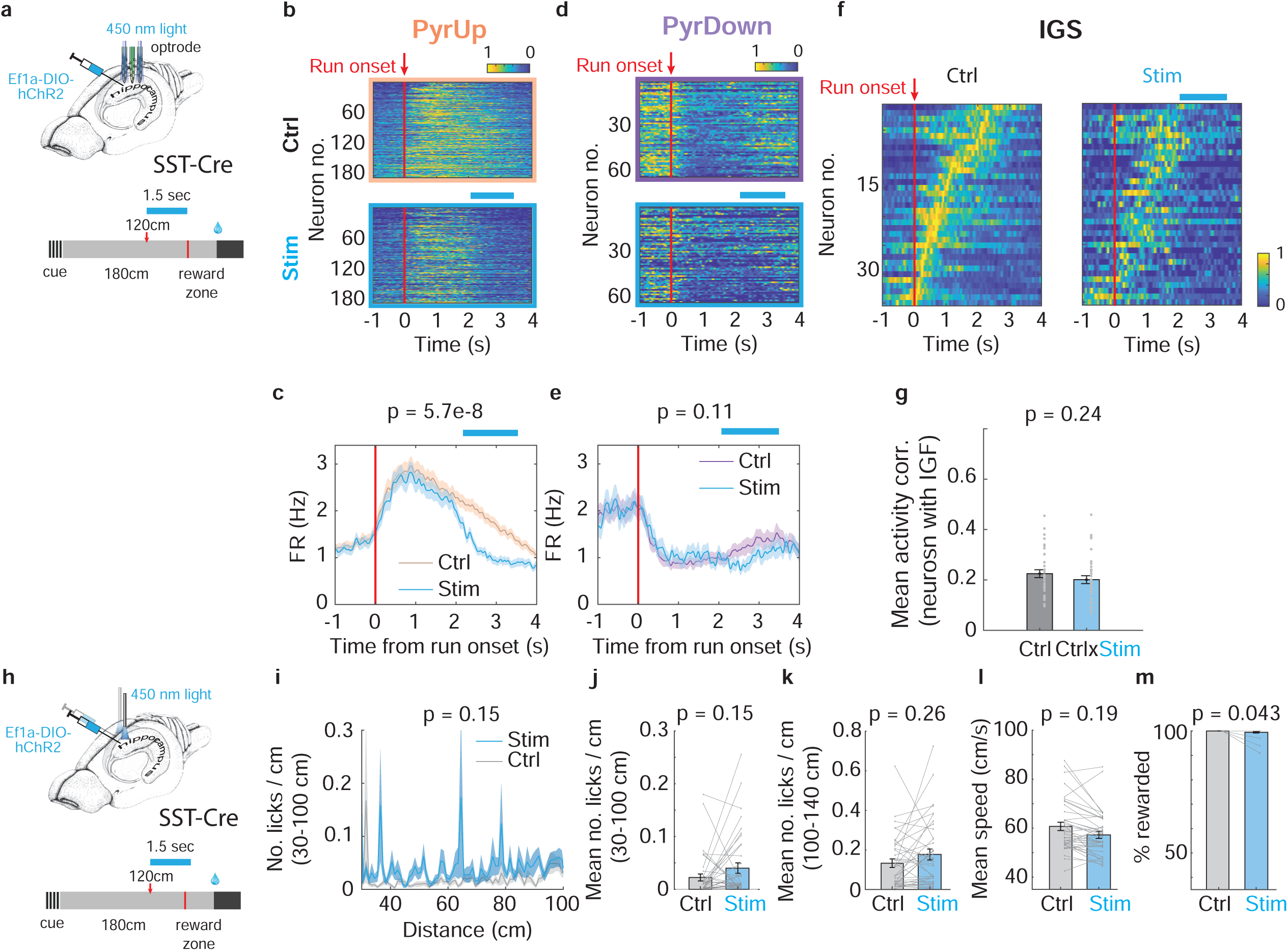
SST interneuron activation in the middle of a trial. (a) SST interneuron activation protocol: Optogenetic stimulation (1.5 s duration, starting at 120 cm during the cue-constant segment) delivered via optrode in a subset of trials (see Methods). (b) Heatmaps of PyrUp neurons during control (top) and stimulation trials (bottom). Data pooled from 3 animals across 8 recordings (183 of 439 neurons). Neurons ordered by firing rate ratio (R = FRaft/FRbef). Blue bar: estimated stimulation window. (c) PyrUp average firing rate profiles for control (orange) and stimulation trials (blue). Wilcoxon rank-sum test (FR 2.5-3.5s): control: 1.70 ± 0.11Hz, stimulation: 0.98 ± 0.08Hz, p = 5.7e-8. (d) Analogous to panel (b), for PyrDown neurons (63 of 439 neurons). (e) Analogous to panel (c), for PyrDown neurons. FR 2.5-3.5s: control: 1.33 ± 0.17Hz, stimulation: 1.03 ± 0.13Hz, p = 0.11. (f) IGS during control (left) and stimulation trials (right). (g) Mean activity correlation among IGF-expressing neurons. Control = 0.22 ± 0.02, stimulation = 0.20 ± 0.02, p = 0.24. (h) Experimental setup to assess the behavioral effects of SST activation. The same protocol as panel (a), in a separate cohort of with bilateral optic fiber implants. (i) Lick histogram from the first 100 cm for control (grey) and stimulation trials (blue). Data from 5 animals, 39 recordings. (j-m) (j) Mean number of licks/cm from the first 100 cm based on panel (i); (k) Mean number of licks/cm between 100-140 cm; (l) Mean running speed; (m) Percentage of rewarded trials for control and stimulation trials.

In contrast, the impact on the PyrDown activity and their rise rates remained insignificant (Fig. 5d-e, rise rates: control: 3.68 ± 0.59s, stimulation: 3.57 ± 0.68s, p = 0.90, two-sample t-test). This lack of effect, despite stimulation occurring during the PyrDown ramping-up period, suggests that EC inputs have a limited contribution to PyrDown dynamics. In addition, our previous findings demonstrated that (1) inactivating SST interneurons after 120 cm did not significantly impact PyrDown activity, likely due to their weak CA3 inputs, and (2) inactivating PV interneurons at run onset, which releases perisomatic inhibition significantly elevated PyrDown activity. Together, these results support the idea that PyrDown neurons might primarily receive perisomatic excitatory inputs that are resistant to dendritic SST modulation.

Again, IGF-expressing neurons showed no significant change in trial-by-trial activity correlations during this manipulation (Fig. 5f-g).

We further assessed the behavioral impact of SST activation by performing bilateral optogenetic activation at 120 cm for 1.5 s in the same cohort of animals as in Fig. 4 (Fig. 5h). This manipulation produced no significant changes in licking, running speed, and a mild effect on the percentage of rewarded trials (Fig. 5i-m, Supplementary Fig. 3c-d). These results contrasted with strong behavioral impairment with PV activation. The preserved performance implies that while PyrUp activity was suppressed, the largely unaffected PyrDown and residual PyrUp activity played a role in maintaining task performance after 120 cm. One possibility is that as animals approached the reward zone, PyrDown activity became stronger, surpassing the declining PyrUp activity around the moment of predictive lick onset (First predictive lick distance: 136.8 ± 6.9cm, first predictive lick time: 3.2 ± 0.2s), and may have exerted greater influence on behavior.

Altogether, we hypothesize that PyrUp dynamics are primarily driven by dendritic inputs (EC/CA3) regulated by SST interneurons, while PyrDown dynamics are mainly shaped by perisomatic inputs gated by PV interneurons.

### A computational model of PyrUp and PyrDown dynamics

To integrate our experimental findings and test the hypothesis on how different excitatory inputs shape PyrUp and PyrDown dynamics under the control of inhibitory interneurons, we developed a biologically informed computational model of the CA1 microcircuit. The model incorporated four neuronal populations^15,28^: pyramidal neurons, PV interneurons, SST interneurons, and intermediate radiatum (RAD) interneurons^21,23,29^. Synaptic connections between these populations were assigned based on defined synaptic strengths and random connectivity probabilities^30^.

CA1 pyramidal neurons were modeled using a two-compartment framework, comprising a dendritic compartment and a somatic compartment^31,32^. The dendritic compartment received simulated distal inputs from EC and proximal inputs from CA3 (Fig. 6a-b). To mimic the inputs driving PyrUp neurons during running, both distal and proximal inputs transiently increased at run onset before decaying as animals approached the reward zone. The somatic compartment integrated dendritic activity with other somatic inputs. To mimic inputs driving PyrDown neurons during reward approaching, somatic inputs gradually ramped toward reward (Fig 6a-b; Eq. 2, Methods). PV, SST, and RAD interneurons were modeled as single-compartment point neurons. PV interneurons synapsed onto pyramidal somata, while SST interneurons inhibited distal dendritic regions and indirectly disinhibited proximal dendrites by suppressing RAD interneurons (Fig. 6a).

**Figure 6.**
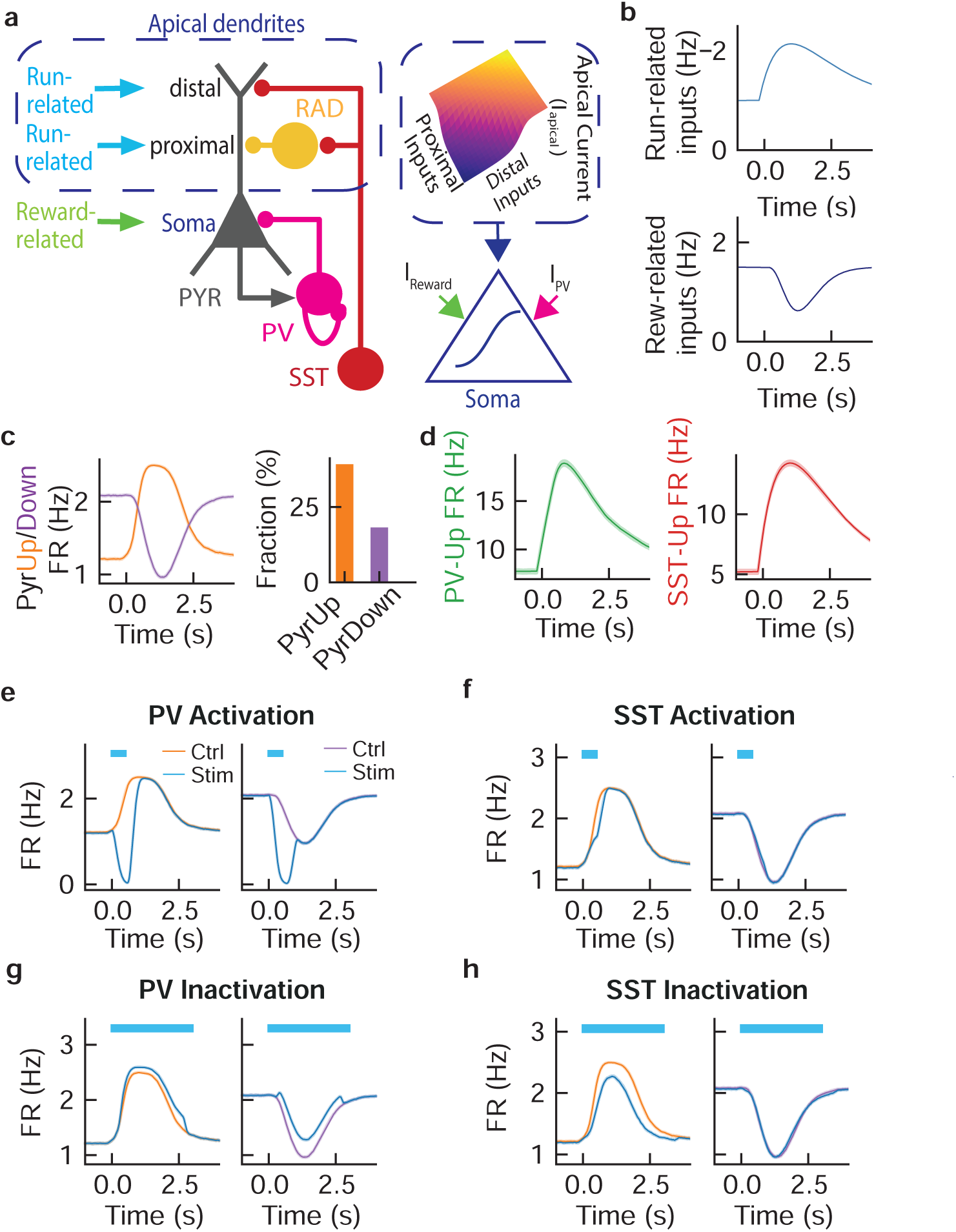
Computational model of CA1 neuronal dynamics during time or distance integration. (a) Model schematic: Network architecture includes four CA1 populations: Pyramidal neurons (n = 2,000) with distinct input compartments: Distal/proximal apical dendrites: Receive run-related inputs (nonlinearly integrated as I_apical_). Perisomatic region: Receives reward-related inputs (I_reward_) and PV-interneuron inputs (I_PV_). Interneurons: Parvalbumin (PV, n = 500), somatostatin (SST, n = 150), and radiatum interneurons (RAD, n = 100). (b) Temporal profiles of run-related inputs (top) and reward-related inputs (bottom), 0 s is run onset. (c) Left: Simulated firing rate dynamics of PyrUp (orange) and PyrDown (purple) neurons. Right: Proportions of PyrUp and PyrDown neurons within the pyramidal population. (d) Firing rate profiles of PV (left) and SST interneurons (right) showing increased activity at run onset. (e) PyrUp (left) and PyrDown (right) dynamics during optogenetic PV activation (blue vs. control) (f) Analogous to panel (e), for SST activation. (g–h) PyrUp/PyrDown dynamics during PV inactivation (g) and SST inactivation (h).

The model replicated key experimental observations. First, it reproduced PyrUp and PyrDown dynamics during the PI task with proportions of neurons in each subpopulation closely matching experimental observations (Fig. 6c). Second, simulated PV and SST interneuron activity mirrored experimental data, where a major population exhibited increased activity at the run onset (Fig. 6d, PV-Up and SST-Up neurons)^14^. Third, optogenetic activation was simulated by injecting positive currents into a fraction of interneurons in either the PV or SST population. These manipulations produced effects consistent with experimental data (Fig. 6e-f). Furthermore, we also replicated our previous results from optogenetic inactivation by injecting a negative current into specific interneuron populations (Fig. 6g-h)^14^.

Overall, the proposed circuit architecture in the model demonstrated the possibility that PyrUp dynamics are primarily shaped by run-related dendritic inputs from CA3 and EC, regulated by SST interneurons, while PyrDown dynamics are preferentially influenced by reward-related perisomatic inputs gated by PV interneurons. By revealing how SST and PV interneurons differentially regulate dendritic and somatic excitation to shape neuronal dynamics, the model provides a mechanistic explanation for the CA1 circuit’s two-phase neural code during time integration.

## DISCUSSION

In our study, we demonstrated two functional subpopulations of CA1 pyramidal neurons—PyrUp and PyrDown neurons—that support a two-phase neural code for encoding time elapsed or distance traveled after a start signal^14^. These two subpopulations exhibit opposite firing dynamics. PyrUp neurons rapidly increase their firing at the onset and then decay, while PyrDown neurons shut down their activity and then ramp up. Their dynamics also inform behavior. The crossover point, where rising PyrDown activity overtakes declining PyrUp activity, consistently precedes predictive licking, suggesting it might serve as a neural marker for imminent reward. Optogenetic experiments further revealed that disrupting the time-encoding phase (phase II), but not the initiation phase (phase I), significantly impairs behavioral accuracy.

Moreover, manipulating PV and SST interneurons provided insights into the circuit mechanisms shaping PyrUp and PyrDown dynamics. Our results support the hypothesis that PyrUp neurons primarily receive locomotion-related dendritic inputs regulated by SST interneurons, while PyrDown neurons mainly receive reward-related perisomatic inputs gated by PV interneurons.

Complementing these experiments, our computational model, incorporating CA1 pyramidal cells and PV/SST interneurons, recapitulated observed neuronal dynamics and optogenetic effects and supported our circuit-level hypothesis, linking dendritic-somatic processing to population-level dynamics.

In summary, these findings offer new insights into how the hippocampal circuits transform neuronal dynamics encoding spatiotemporal information into predictive behavior.

### PyrUp and PyrDown activity together provides a neural signal of imminent reward

Our results demonstrate that the complementary dynamics of PyrUp and PyrDown neurons may serve as a neural predictor of reward. Specifically, the population-level crossover point, where rising PyrDown activity surpasses decaying PyrUp activity, consistently occurs just before the onset of predictive licking. This alignment persists even in trials where the animal did not accurately integrate time or distance, suggesting that the crossover point acts as a robust neural marker of imminent reward, independent of behavioral accuracy.

Why might this crossover point signal the onset of predictive licking? Clues lie in the functional and circuit-level properties of these two subpopulations. First, PyrUp neurons exhibited synchronized activation at run onset, followed by gradual decay toward reward, reflecting dominant excitation during running. Conversely, the PyrDown neurons displayed shutting down activity at run onset and progressively ramping up toward reward, suggesting dominant excitation during reward approaching. Second, PyrUp neurons preferentially encoded run-related events (e.g. run onset), while PyrDown neurons biased towards reward-related events (e.g., predictive licking, reward delivery). Third, the PV and SST manipulation results support that PyrUp neurons mainly process dendritic inputs from EC and CA3, whereas PyrDown neurons preferentially receive somatic inputs.

Together, our results support that PyrUp and PyrDown neurons complement each other: PyrUp neurons process run-related spatial-temporal signals, while PyrDown neurons encode reward-relevant information. As run-related excitation driving PyrUp activity wanes and reward-related excitation driving PyrDown activity strengthens, the crossover point merges as an imminent reward signal. Therefore, PyrUp and PyrDown dynamics not only track elapsed time and distance but also play a role in linking hippocampal computation to predictive behavior, bridging memory encoding and goal-directed action.

### PyrDown neurons preferentially receive reward-related perisomatic inputs regulated by PV interneurons

Our previous work suggested that PV interneurons preferentially modulate PyrDown neurons. Two observations supported this hypothesis. First, PyrDown activity started to decrease at run onset only after the initial rise in PV interneuron firing, while PyrDown activity reached its minimum shortly after PV activity peaked^14^. Second, optogenetic PV inactivation at run onset selectively impaired PyrDown activity by preventing their initial shutdown without significantly affecting PyrUp neurons^14^.

A plausible explanation for these results might involve preferential synaptic connectivity between PV interneurons and PyrDown neurons. However, several lines of evidence challenged this possibility. First, while previous studies^33,34^ report differential PV connectivity with deep versus superficial CA1 pyramidal neurons, PyrUp and PyrDown neurons showed no significant spatial segregation within the pyramidal layer^14^. This result suggests that both subpopulations comprise a mix of genetically distinct deep and superficial neurons. Second, cross-correlogram analysis^27^ detected no significant differences in monosynaptic connectivity from PV interneurons to PyrUp versus PyrDown neurons (PV to PyrUp: 0.008±0.003, PV to PyrDown: 0.017±0.005, p = 0.49). However, this method may underestimate connectivity compared to paired recordings^33^. Third, PV activation non-selectively suppressed PyrUp and PyrDown activity, implying that PV interneurons do not exclusively modulate PyrDown neurons.

These inconsistencies led us to propose an alternative hypothesis: PyrDown neurons receive dominant perisomatic excitation from reward-related inputs that are regulated by PV interneurons. This hypothesis aligns with three observations: First, PyrDown activity ramped up during reward approaching, consistent with reward-related excitation. Second, PV interneurons preferentially inhibit perisomatic regions, and PV inactivation primarily impaired PyrDown activity, suggesting PV-mediated regulation of PyrDown activity^14^. Third, SST interneuron activation and inactivation, which mainly influence dendritic inputs from EC and CA3, left PyrDown neurons largely intact^14^. This result suggests dendritic inputs have a limited contribution to PyrDown dynamics. We also validate this hypothesis using a computational model to simulate interactions between PV activity, reward-related inputs, and CA1 pyramidal neuron dynamics. The model demonstrated how PV-mediated perisomatic inhibition gates reward-related inputs to shape PyrDown dynamics, providing mechanistic evidence for compartment-specific processing in CA1.

### The role of PV interneurons in regulating time or distance integration

Our previous work demonstrated that PV interneuron inactivation at run onset led to behavioral deficits related to the initiation of time or distance integration^14^. This finding supports that a rapid increase in PV activity at run onset is necessary to suppress PyrDown activity and start the integration process.

In contrast, the current study revealed that PV activation, which transiently silenced pyramidal neurons at run onset, did not impair task performance. This result has two possible interpretations. First, during phase I, the precise firing pattern of CA1 pyramidal neurons may not be critical for supporting accurate integration of time or distance, as this phase primarily signals the start of integration rather than encoding elapsed time or distance. Second, PV activation enhances the “start” signal by further suppressing PyrDown neurons. However, the behavioral performance in control trials was already near optimal, potential enhancement was not detectable.

On the other hand, PV activation in the middle of a trial, corresponding to phase II, significantly impaired task performance, underscoring the necessity of intact PyrUp/PyrDown ramping dynamics for integration accuracy. However, this result may bear an alternative explanation: the PV-mediated rapid shutdown of PyrDown neurons prematurely triggered a new initiation signal, causing the animal to overshoot the reward zone. Yet, our behavioral data did not show acceleration in running or reduction in licking during PV activation, and neuronal data did not reveal a clear new initiation phase following stimulation. These observations indicate that the performance deficits likely reflect disrupted integration accuracy rather than reinitiation of integration. Together, these results support that PV activity is necessary for initiating integration but insufficient to fully control it.

Mechanistically, PV interneurons can dynamically regulate PyrDown activity in two phases. At run onset, rapidly increased PV activity suppresses PyrDown neurons, thus signaling the start of integration. During reward approach, declining PV activity gradually disinhibits reward-related inputs, permitting PyrDown activity to ramp up. Thus, PV interneurons appear to modulate both phases of PyrDown dynamics, orchestrating integration initiation via perisomatic inhibition and reward anticipation through progressive disinhibition.

### PyrUp neurons preferentially receive run-related dendritic inputs regulated by SST interneurons

Our findings, spanning prior^14^ and current work, establish that somatostatin-positive (SST) interneurons predominantly modulate PyrUp neurons. Two lines of experimental evidence support this conclusion. First, SST inactivation at run onset primarily suppressed PyrUp activity without significantly affecting PyrDown activity^14^. Second, SST activation similarly exerted preferential suppression on PyrUp neurons. Cross-correlogram analysis^27^ revealed no significant differences in monosynaptic connectivity from SST interneurons to PyrUp versus PyrDown neurons, suggesting that connectivity biases alone may not explain these effects.

We propose an alternative mechanism: PyrUp neurons preferentially receive excitation during locomotion, driven predominantly by dendritic EC and CA3 inputs, both regulated by SST interneurons. SST interneurons regulate these inputs through dual mechanisms: direct inhibition of distal EC inputs, and indirect disinhibition of proximal CA3 inputs via suppression of intermediate interneurons. This hypothesis aligns with two observations. First, SST inactivation at run onset suppressed PyrUp activity potentially by indirectly inhibiting CA3 inputs, suggesting a contribution of CA3 inputs in shaping PyrUp dynamics. Second, SST activation reduced PyrUp activity primarily through direct inhibition of EC inputs. These experimental observations were further validated by simulations in our computational model.

Together, these results position SST interneurons as regulators of distinct excitatory afferent pathways, selectively shaping PyrUp activity to control time or distance integration during running.

### The role of SST interneurons in regulating time or distance integration

Our previous work showed that SST interneuron inactivation after run onset impaired behavioral accuracy in the PI task^14^. This finding supports our hypothesis that elevated SST interneuron activity favors the processing of CA3 inputs that carry spatiotemporal information. By disinhibiting CA3 through suppressing intermediate interneurons, SST interneurons establish an integration window that facilitates PyrUp activity and supports the accuracy of time or distance integration.

In the current study, SST activation revealed the second regulatory role of these interneurons: direct inhibition of EC inputs. However, SST activation during the time-encoding phase (phase II) produced no significant behavioral changes. This lack of effect suggests two non-exclusive mechanisms. First, PyrDown activity, which was largely unaffected by SST activation, may help maintain integration accuracy. Second, SST activation indirectly disinhibits CA3 inputs and partially counterbalances the suppression of EC inputs. The latter possibility is supported by the partial suppression of PyrUp activity and a mild reduction in their decay rates during stimulation.

These findings collectively reveal that SST interneurons dynamically balance EC and CA3 excitatory inputs, thereby preferentially regulating PyrUp dynamics and supporting time or distance integration.

### Computational model of the PyrUp and PyrDown dynamics

Our computational model incorporates PV and SST interneurons, which exert compartment-specific control over pyramidal neurons. PV interneurons target somatic regions to regulate global firing output, while SST interneurons modulate dendritic integration by inhibiting distal EC inputs and disinhibiting proximal CA3 pathways. The model assumes spatially segregated excitatory drives: run-related inputs (simulating CA3 and EC afferents) target proximal and distal dendrites, respectively, while reward-related inputs converge on perisomatic regions. This circuit architecture successfully recapitulates experimentally observed neuronal dynamics and optogenetic perturbation outcomes. The model establishes a unified circuit framework for understanding the two-phase neural code and offers new insights into how local hippocampal circuits transform spatiotemporal inputs into memory-guided predictive behavior.

## ACKNOWLEDGMENTS

We thank D. Fitzpatrick for discussions; M. Klement, N. Daniel and machine shop at MPFI for making mechanical parts for the experimental setups; J. Wells and ARC at MPFI for taking care of animals. This work was funded by Max Planck Society and Max Planck Foundation, NIH R01 NS119503.

## AUTHOR CONTRIBUTIONS

Y.W. conceived the project. Y.W. and R.H. designed experiments (with input from D.P.). R.H. performed electrophysiological experiments. D.P., Y.W, and X.Z. performed behavioral experiments with optogenetics. Y.W. and R.H. analyzed data. C.P., A.R. and Y.W. constructed the computational model, Y.W. wrote the manuscript, with contribution from all the authors.

---

**Supplementary Figure 1.**
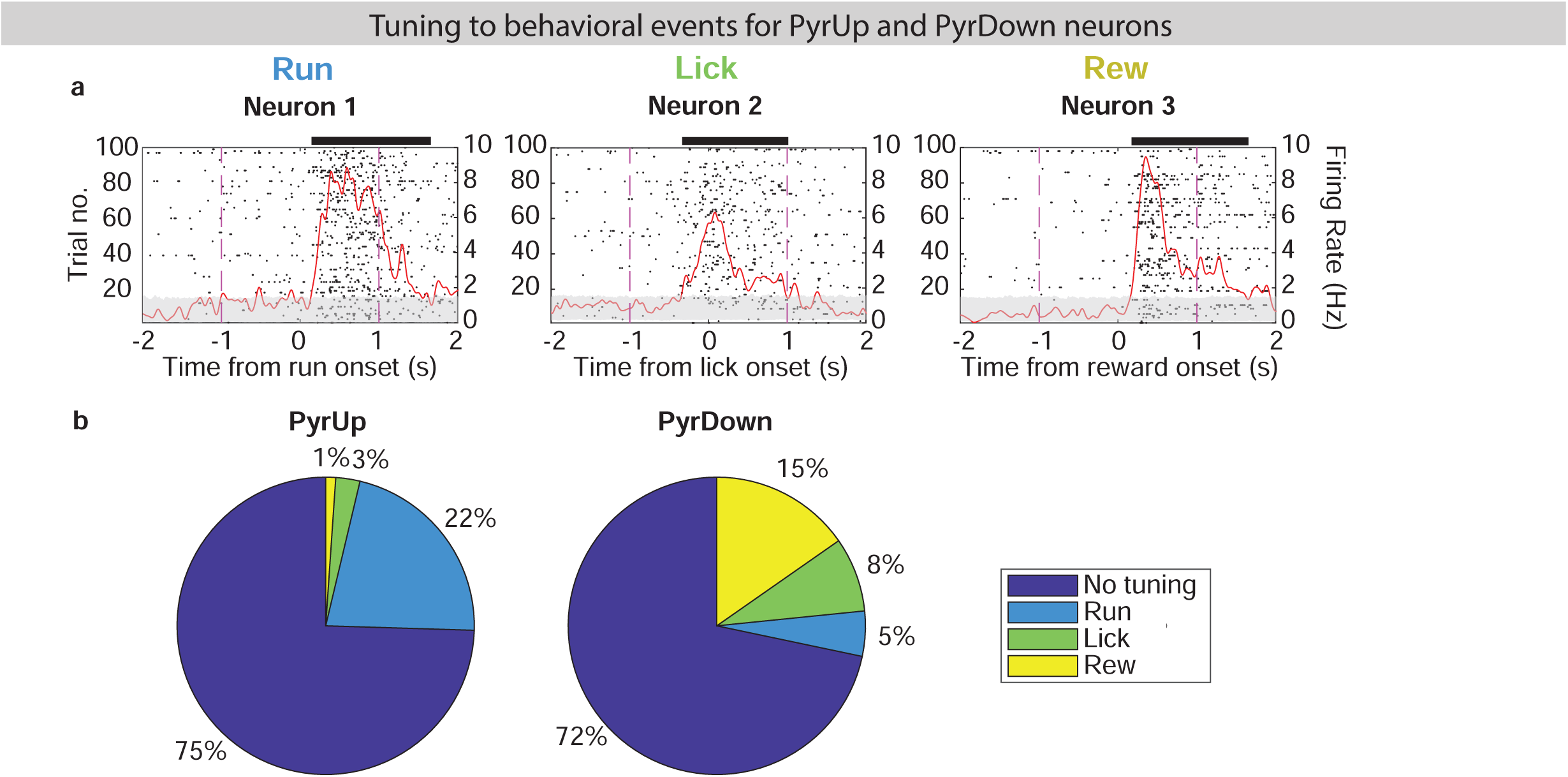
Run-related encoding in PyrUp dynamics and reward-related encoding in PyrDown dynamics, related to Figure 1. (a) Example raster plots and firing rate profiles (red lines) for pyramidal neurons significantly tuned to: Run onset (left), First lick (middle), and Reward delivery (right). Gray shading: 1st–99th percentile of shuffled data. Black horizontal bars: Time windows of significant tuning. (b) Proportions of PyrUp (left) and PyrDown (right) neurons classified by tuning to behavioral events: No Peak: No significant tuning, Run: Maximal tuning at run onset, Lick: Maximal tuning at first lick, Rew: Maximal tuning at reward delivery. Neurons with multi-event tuning were exclusively categorized based on the tuning with the highest firing rate modulation amplitude.

**Supplementary Figure 2.**
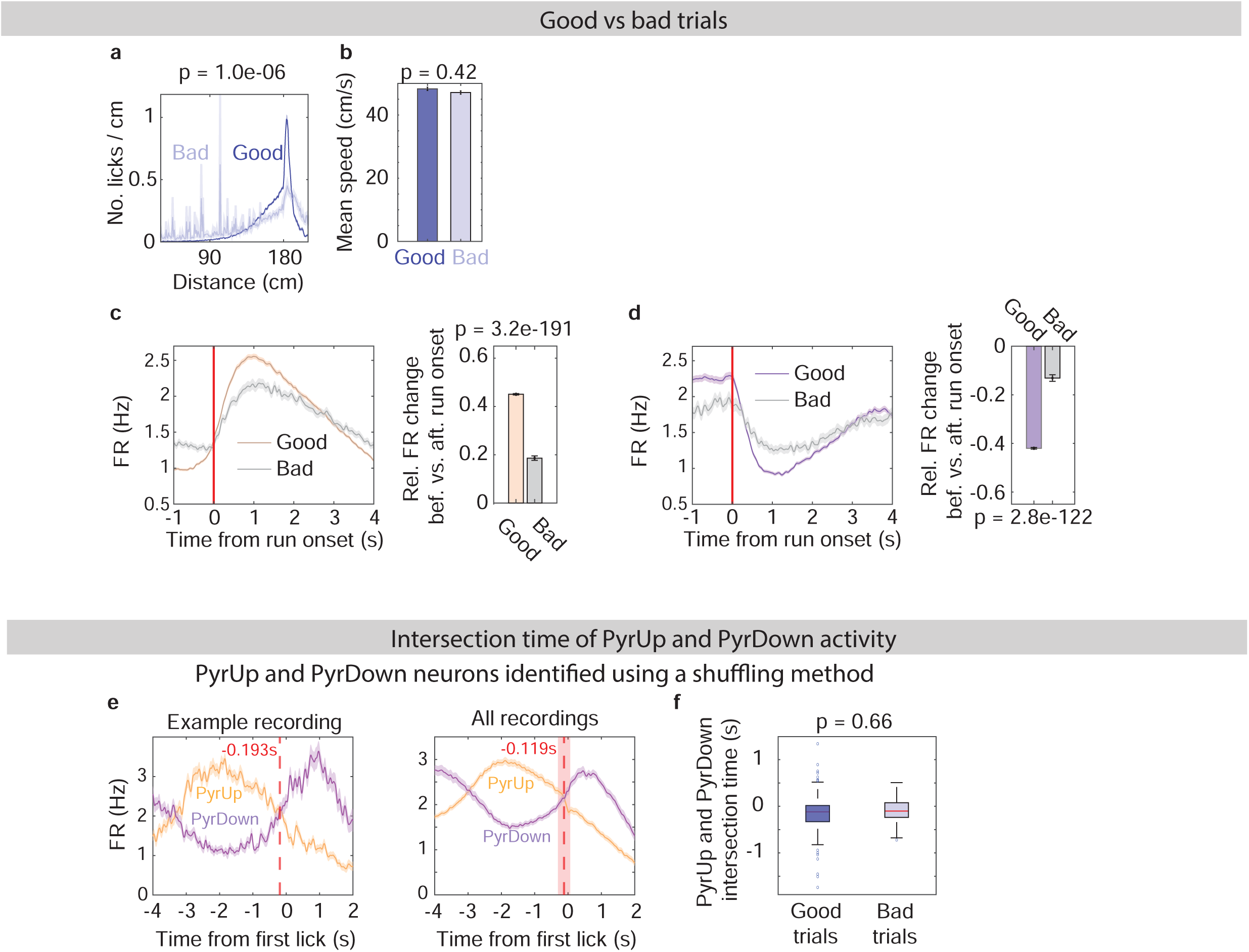
Behavioral relevance of PyrUp and PyrDown activity, related to Figure 1. (a-b) Lick histogram (reported p-value is for mean number of licks/cm between 30-100 cm) (a) and mean running speed (b), for good (blue) and bad trials (light blue). (c) Left: Average firing rate profiles (±SEM) of PyrUp neurons in good vs. bad trials (left). Right: Relative firing rate change before vs. after run onset, calculated as (FRbef – FRaft)/(FRbef + FRaft) (right). (d) Analogous to panel (c), for PyrDown neurons. (e) Left: first-predictive-lick aligned firing rate profile averaged across all PyrUp (orange) and PyrDown (purple) neurons from one example recording. Red dotted line: their intersection. Right: Aggregated data from all the recordings. Median intersection time (red dotted line) ± MAD (shading). PyrUp and PyrDown neurons are identified as neurons with significant R*-*value deviations from shuffled data. (f) Intersection time distributions in good versus bad trials. Neurons identified as in panel (e).

**Supplementary Figure 3.**
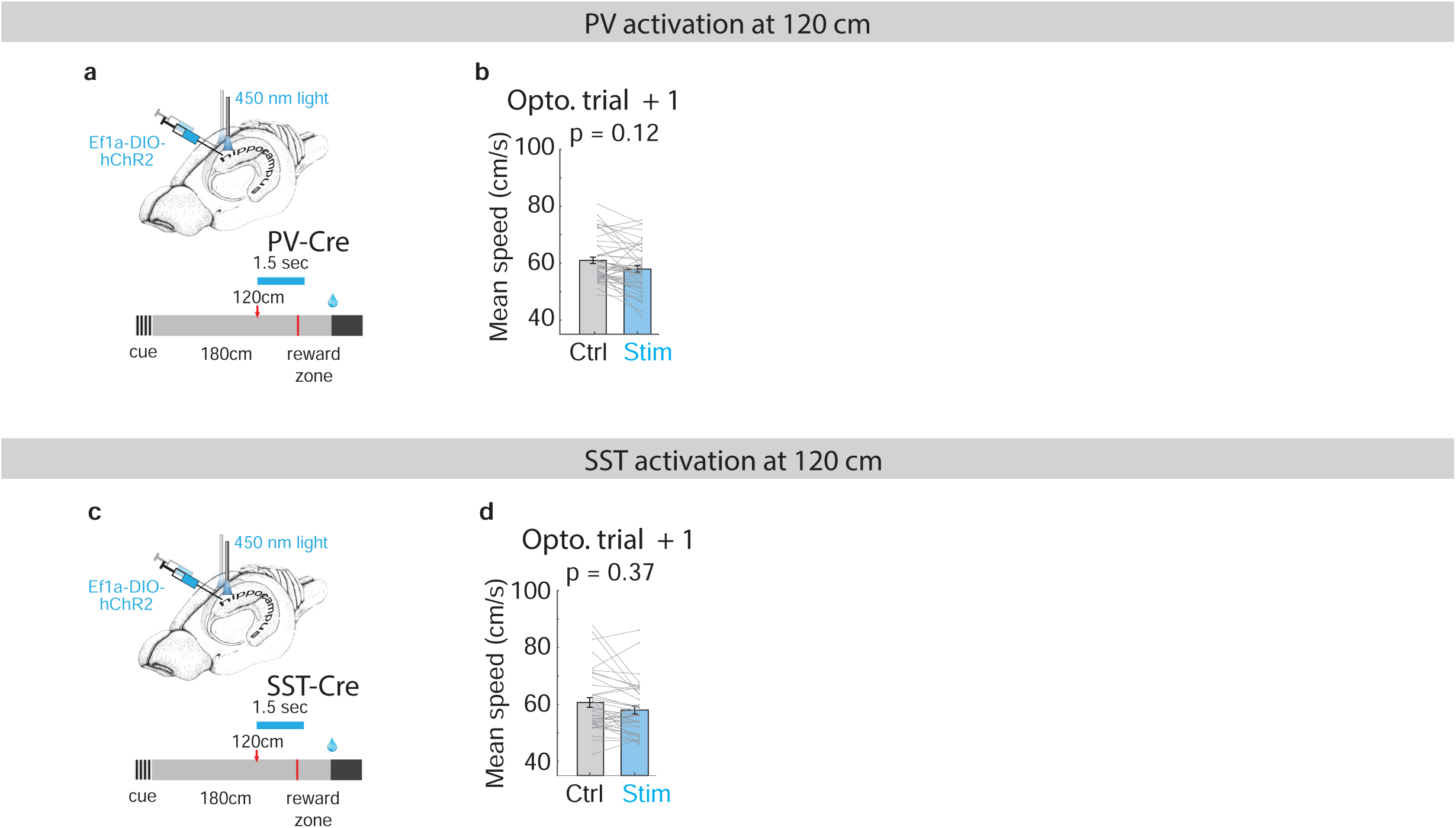
Behavior during PV and SST activation in the middle of the trial, related to Figures 3 and 5. (a) Experimental setup to assess the behavioral effects of PV activation in the middle of the trial, as in Fig. 3h. (b) Mean running speed of trials immediately following the PV activation trials. (c-d) Analogous to panels (a-b), for SST activation.

**Supplementary Figure 4.**
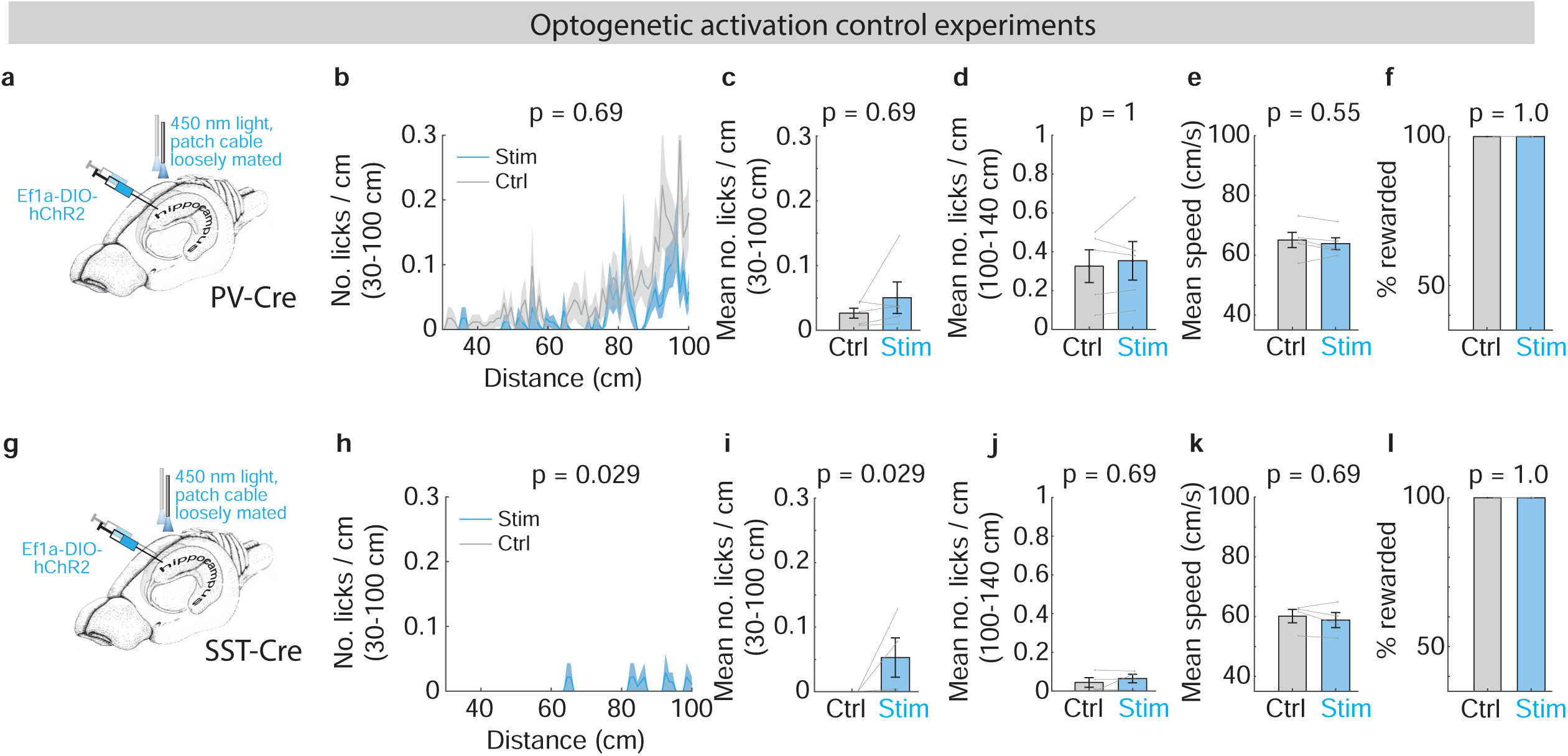
Control experiments for bilateral optogenetic activation, related to Figures 2 to 5. (a-f) Control experiments for optogenetic activation of PV interneurons (2 animals). (a) Experimental design. Light delivery protocols are identical to PV activation experiments, but the patch cables were adjusted not to be contact the implanted fiber optic cannulas, precluding light delivery into the brain. (b) Lick histogram from the first 100 cm for control (grey) and stimulation trials (blue). (c) Mean number of licks/cm from the first 100 cm based on panel (b); (d) Mean number of licks/cm between 100-140 cm; (e) Mean running speed; (f) Percentage of rewarded trials for control and stimulation trials. (g-l) Analogous to panels (a–f), for optogenetic activation of SST interneurons (2 animals).

**Supplementary Figure 5.**
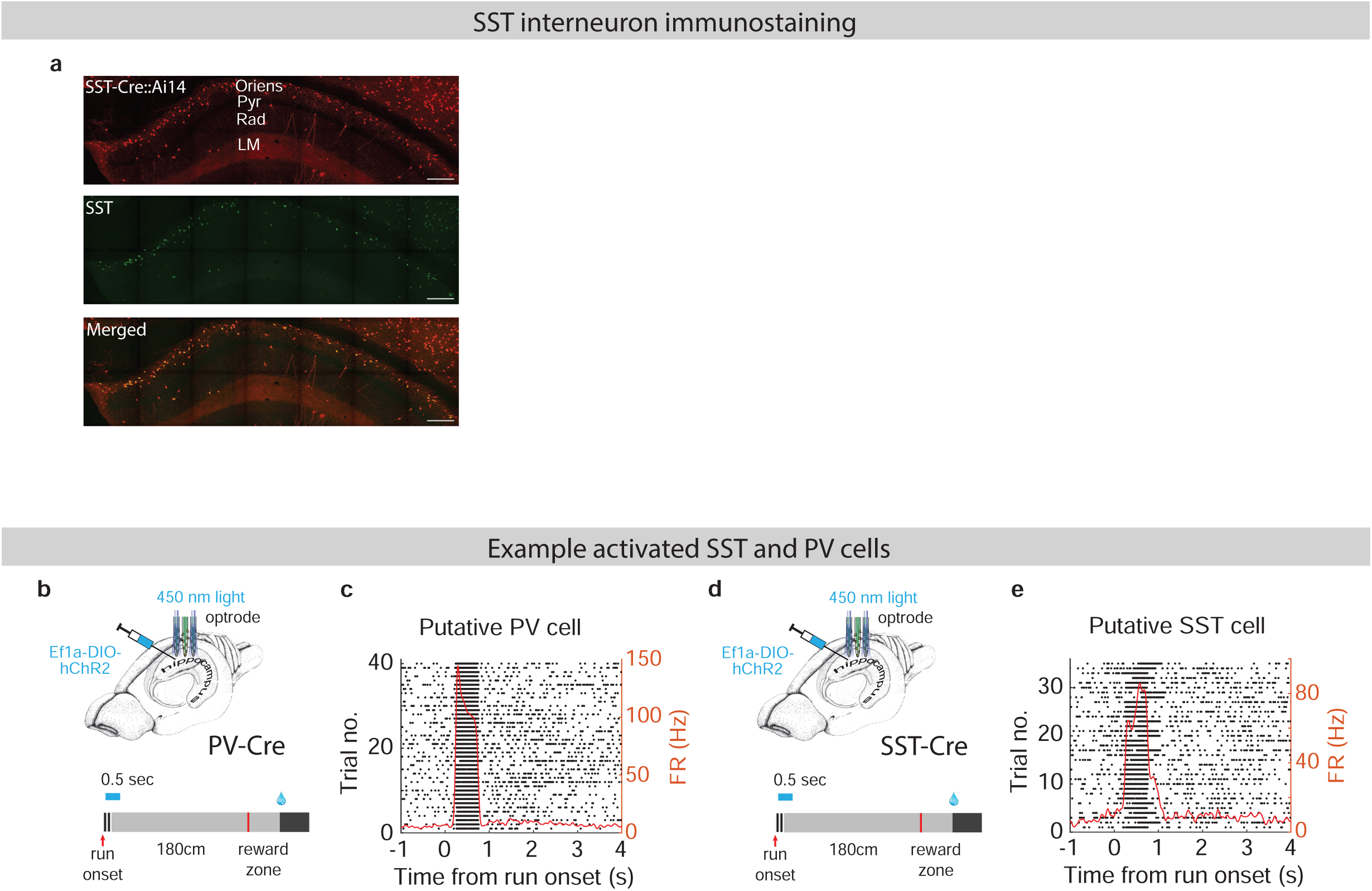
SST and PV interneurons, related to Figures 2 and 4. (a) Confocal images of CA1 from an SST-CrexAi14 mouse. From top to bottom, tdTomato expression, SST immunostaining (GFP), and merged image. Scale bar: 200 um. (b) Experimental setup to assess the behavioral effects of PV activation at run onset, as in Fig. 2h. (c) Spike raster plot and average firing rate profile (red line) of an example putative PV interneuron that is effectively activated by optogenetic stimulation. (d-e) Analogous to panels (b-c), for SST activation.

## METHODS

### EXPERIMENTAL MODEL AND SUBJECT DETAILS

#### Mice

This study was based on both male and female mice (age > 8 weeks). Male mice were preferentially used in the running tasks because they were found to exhibit more consistent running behavior. We used three mouse lines: C57Bl/6J (JAX #000664), PV-IRES-Cre (JAX #017320), and SST-IRES-Cre (JAX #013044)^35^, and Ai14 (JAX # 007914)^36^. All procedures were in accordance with protocols approved by the Institutional Animal Care and Use Committee at Max Planck Florida Institute for Neuroscience. Mice were housed in a 12:12 reverse light: dark cycle and behaviorally tested during the dark phase.

### METHOD DETAILS

#### Virtual reality setups

For the virtual reality (VR) setups, a small treadmill (Janelia Design) was positioned in front of two visual displays, such that after head-fixation, the eyes aligned with the two visual displays. Customized software was written using Psychtoolbox to display the virtual environment. An Arduino control board was used to control when to display the visual stimuli. For the small treadmill, the animal’s speed was measured using an encoder attached to the back wheel axis. A microprocessor-based (Arduino) behavioral control system (the miniBCS board, designed at Janelia) interfaced with a MATLAB graphical user interface controlled the trial structure, the water valve, and the encoder. A separate lick port detector (designed at Janelia) was used to convert a touch on a metal lick port into a digital pulse and to send the information to the miniBCS board. In another version of the VR setup, instead of using two monitors, a curved screen was used where the virtual reality environment was projected onto the curved screen using a projector (AKASO Mini Projector, a design kindly shared by Christopher Harvey’s lab at Harvard University). In this case, customized software written with Unity was used to display the virtual environment.

In addition, a separate microprocessor (Arduino) interfaced with a customized MATLAB graphical user interface was used to operate the laser or laser diode on-off for the optogenetic perturbation experiments and to control the closed-loop manipulation of PV and SST interneurons. Behavioral data were monitored and recorded as well using MATLAB functions.

#### Behavioral training

Before any surgery was performed, running wheels were added to the home cages. At 3–5 days after the headbar or fiber implantation, mice were placed on water restriction (typically 0.8-1 ml per day). After each training or recording session, mice were supplemented with additional water to ensure the amount of water intake per day. After habituating the animals to the treadmill for at least 1-2 days, animals were trained to run head-fixed on the treadmill.

In the PI task, a trial started with a visual grating stimulus that lasted either 0.5s or 1s, and stayed the same within each session. After that, a grey-colored stimulus turned on. The animal was required to run for a fixed distance (180 cm) while this visual stimulus stayed constant, and then it must lick within a 40 cm (in some sessions, it is 80 cm) unmarked reward zone to receive a water reward. The first lick in the reward zone triggered a reward, and simultaneously the screens turned black for either 0.5s or 1s (kept constant within each session). If no lick occurred in the reward zone, the screens turned black to signify the error for 0.5-2s (kept constant within each session). It took about 2 weeks for the animals to learn this task.

#### Virus injection, headbar, cannula and fiber implantation

Adult mice (2-4 months of age) underwent aseptic stereotaxic surgery to implant a custom lightweight 3D printed headbar under isoflurane anesthesia (2-3% for induction, 1%–1.5% for maintenance). Buprenorphine SR LAB (0.5 mg/kg, SC) or buprenorphine (0.1 mg/kg, SC), and Meloxicam SR (5 mg/kg, SC) were administered immediately after the surgery. For acute extracellular recordings, after training to perform behavior tasks, craniotomies were performed which were centered around anteroposterior 2.1 mm from bregma, mediolateral +/-1.7 mm from the midline to target dorsal CA1 region. Meanwhile, a ground wire was attached to the headbar, and a thin layer of silver paint (GC ELECTRONICS 22-0023-0000) was applied to the surface of the headbar for noise reduction during the recording. The skull was covered using KwikSil (World Precision Instruments) and was only removed during recordings.

For extracellular recordings with optogenetic manipulation of interneurons, AAV5-Ef1a-double-floxed-hChR2(H134R)-mCherry-WPRE-hGHpA (Addgene 20297, titer 1.4e+13) was used for optogenetic activation. 50 nl virus per hemisphere was injected bilaterally in the CA1 of PV-IRES-Cre and SST-IRES-Cre mice. The injection was done at least 3 days before the headbar implantation, and at least 3 weeks before the recording. The following coordinates were used for viral injections: 2.1 mm from bregma, mediolateral +/-1.7 mm from the midline, and 1.24 mm dorsoventral from the brain surface. The injection system comprised a pulled glass pipette (broken and beveled to 15-20 μm inside diameter; Drummond, 3-000-203-G/X), backfilled with mineral oil (Sigma). A fitted plunger was inserted into the pipette and advanced to displace the contents using a manipulator (Drummond, Nanoject II or Nanoject III). Retraction of the plunger was used to load the pipette with the virus. The injection pipette was positioned onto a Kopf manipulator.

For optogenetic experiments without extracellular recordings, the viral injection procedure was the same as described in the last paragraph. At least 3 days after the viral injection, optical fibers (core diameter of 200 um) were chronically implanted bilaterally to target the CA1 region using the following coordinates: 2.1 mm from bregma, mediolateral +/-1.7 mm from the midline and 0.9 mm dorsoventral from the brain surface.

#### Optogenetics without extracellular recordings

For each optogenetic session, a control session (∼40 trials) was performed. It was then followed by a stimulation session (∼100 trials in total), where the photostimulation was deployed every third trial. After that, there was a post-stimulation control session (∼40 trials). To prevent mice from distinguishing photostimulation trials from control trials through stimulation light, a masking blue light (470 nm LEDs (Thorlab)) was on throughout the sessions. For ChR2, we used a 473 nm laser (Ningbo Lasever Inc.). The laser power used was <= 5 mW at the tip of the fiber.

For interneuron activation, the stimulation lasted 0.5 or 1.5 seconds. The stimulation can be triggered at (1) the running onset or (2) 120 cm location during the cue-constant segment. The run onset stimulation was triggered at the first time point where the animal’s speed exceeded 10 cm/s for at least 200 msec after the trial start. The stimulation trial types were randomly selected for each stimulation session. The behavior analysis was performed using all the trials during the stimulation session. Only considering the stimulated trials led to similar results.

As a control, in some recording sessions, the optogenetics patch cables were only loosely mated with the implanted fiber optic cannulas, preventing light from propagating into CA1.

#### Acute extracellular electrophysiology

Two days before the recording, the animal was acclimated to the recording condition: including turning on the microscope light, removing Kwiksil that covered the skull, and adding saline to keep the craniotomy wet. On the day before the recording, craniotomy was opened. During electrophysiological recordings, a 64-channel silicon probe (neuronexus, buzsaki64sp) was slowly lowered into the hippocampal CA1 region. Data from all channels were filtered (0.3 Hz to 10 kHz), amplified (gain = 400), and continuously sampled at 20 kHz using the Amplipex system (16-bit resolution)^37^. Time stamps of behavioral events and electrophysiological recording data were synchronized, recorded, and stored on a computer.

For extracellular recordings with optogenetics, a 64-channel silicon probe with fibers mounted on the shanks (neuronexus, buzsaki64sp-OA64LP) was used. Photostimulation followed the same protocol as described in the optogenetics section above. The light source was a self-constructed laser diode array (6 diodes, 450 nm, osram-os, PL450B). Each diode was coupled to one shank on the probe. 2 to 5 diodes were coactivated during the stimulation. The stimulation laser power was < 1.2 mW at the tip of the probe.

#### Spike sorting

To identify spikes, we performed off-line spike sorting on the recorded files following published methods using Klusters^38^ and Kilosort^39^. Units were further selected based on the percentage of spikes that violated refractory period and the mahalanobis distance from other units.

#### Histology

Mice were perfused transcardially with PBS followed by 4% PFA. Brains were post-fixed overnight and transferred to PBS before sectioning using a vibratome. Coronal 50 μm free-floating sections were processed using standard fluorescent immunohistochemical techniques. PV immunoreactivity (Swant, GP72; 1:2000, guinea pig). The primary antibody was visualized by Alexa Fluor 488 AffiniPure Donkey Anti-Guinea Pig IgG (H+L) (1:1,000, Jackson ImmunoResearch Laboratories, 706-545-148). SST immunoreactivity (Millipore, MAB354; 1:250, rat). The primary antibody was visualized by Alexa Fluor 488 AffiniPure Donkey Anti-Rat IgG (H+L) (1:1,000, Jackson ImmunoResearch Laboratories, 712-545-153).

### QUANTIFICATION AND STATISTICAL ANALYSIS

#### Behavior analysis

To calculate how the speed changed over distance during the cue-constant segment, the histogram of speed within each trial was calculated using 1 mm distance bins. 15 trials were randomly selected from each session, and the average and SEM of speed were calculated across all the selected trials from all the sessions in each task. Similarly, to calculate how the number of licks changed over distance, a lick histogram with a bin size of 1 cm was first calculated for each trial. Then the same calculation as the speed was applied to get the average and SEM. To calculate the overall mean speed or lick within a certain distance range, the mean speed or lick number was first calculated for each session, then averaged across sessions from the same task. To calculate total stop time, we first smoothed running speed using a moving average with an 80ms window, and then summed the time when the speed was lower than 1 cm/s from the running onset to the end of the trial.

#### Extracellular recording analysis

##### Alignment with sensory or motor cues

For the PI tasks, both the neuronal activity and the behavior of each trial were aligned with either the start cue onset or the running onset. The running onset for each trial was defined as the onset of the first running bout that lasted more than 0.3 sec with a speed > 10cm/s, starting from the previous trial after the animal had traveled the 180 cm cue-constant segment. The onset was when the running speed reached 1 cm/s before the running bout and after 180 cm in the cue-constant segment of the previous trial. If the speed never reached 1 cm/s, the time point when the speed was the lowest was considered as the onset, and this trial was classified as “no clear run onset”.

##### Firing rate profile of each neuron

Pyramidal neurons were identified based on their spike waveforms, and if their firing rates fell between [0.15 7] Hz. The spike train of each pyramidal neuron was first smoothed with a Gaussian function (s.d. = 30 ms) and then averaged across trials to get the average firing rate profile of that neuron. When examining how the firing rate changes over distance, the spike train was first binned using 1 mm spatial bins, and then smoothed with a Gaussian function (s.d. = 20mm). Normalized firing rate profiles of PyrUp or PyrDown neurons were calculated by first normalizing the firing rate profile of each neuron by its max firing rate and then averaging across the selected population.

##### Firing field identification

The average firing rate profile of each neuron was used to identify firing fields (internally generated fields). To identify firing fields over distance, the following criteria were applied: 1) a minimum mean firing rate of 0.09 Hz; 2) minimum mean trial-by-trial Spearman correlation of the firing rate profile > 0.15 and minimum spatial information^40^ > 0.25 bits/s; or minimum mean trial by trial correlation > 0.09 and minimum spatial information > 0.7 bits/s. 3) total number of trials > 15, and the neuron was active in at least 40% of the trials. 4) the maximum field width was 150 cm, measured by the width when the firing rate fell below 10% on both sides of the peak firing rate. 5) there was no other peak within 30 cm outside the field. The parameters were chosen based on visual inspection.

To identify firing fields over time, we used similar criteria as mentioned above. Some of the parameters were readjusted based on visual inspection: 1) minimum mean trial-by-trial correlation > 0.12 and minimum temporal information > 1 bits/s; or minimum mean trial-by-trial correlation > 0.09 and minimum temporal information > 2.5 bits/s. 2) The maximum field width was set to 4.5 sec. 3) There was no other peak within 1.7 sec outside the field.

##### Criteria for good and bad trials

The criteria for identifying good trials in the PI task are: 1) The animal came to a complete stop at the reward location of the previous trial just before run onset (speed < 1 cm/s). 2) The subsequent running was mostly continuous, with a total stop time shorter than 2s after run onset. 3) The animal obtained a reward. Otherwise, the trial was considered a bad trial.

##### Identification of PyrUp and PyrDown neurons

The change in firing rate around the run onset for each neuron was defined as the ratio between the average firing rate in a window 0.5 to 1.5 seconds (FRaft) and −1.5 to −0.5 seconds (FRbef) about the running onset (FRaft/FRbef). The PyrUp neurons were those whose FRaft/FRbef > 3/2, while the PyrDown neurons were those whose FRaft/FRbef < 2/3. PyrOther neurons were those whose FRaft/FRbef were between 3/2 and 2/3. The relative firing rate change of the average firing rate profile around run bouts and trial run onsets was calculated for each neuron in a window 0.5 to 1.5 and −1.5 to −0.5 seconds about the running onset with the formula (FRbef – FRaft)/(FRbef + FRaft).

The same criteria were used to identify PV-Up and SST-Up neurons.

To avoid bias in grouping neurons, an additional shuffling method was used to identify PyrUp and PyrDown neurons. Shuffled distributions of FRaft/FRbef were created by performing randomized circular shifts to the firing rate profile of each trial and recalculating FRaft/FRbef 1000 times. Neurons with a firing rate change above the 95th percentile or below the 5th percentile of the shuffled distribution were labeled as PyrUp or PyrDown neurons, respectively.

##### Decay/rise time constant of PyrUp/PyrDown dynamics

For each PyrUp neuron, the first peak in its mean firing rate profile that exceeded the 99 percentile of the shuffled mean firing rate profile (shuffled 500 times) was identified. The decay time constant was extracted by fitting the post-peak mean firing rate profile with the function a*exp(bx), where the time constant is defined as −1/b.

Similarly, for each PyrDown neuron, the significant peak 1 sec after run onset was identified as for PyrUp neurons. The time constant was calculated by fitting the function a*exp(bx) from the peak backward to 1 sec after run onset.

##### Crossover point of population-averaged PyrUp and PyrDown activity

For each trial, neuronal activity was aligned to the time of the first predictive lick. The activity was then averaged separately across PyrUp and PyrDown neurons identified in the recording session. The crossover point (i.e., the time when PyrUp and PyrDown activity profiles intersected) was determined either (1) individually for single trials or (2) after averaging the PyrUp and PyrDown activity across all trials. Finally, we calculated the median and median absolute deviation (MAD) of crossover times across all recording sessions.

##### Tuning to behavioral events

Average firing rate curves aligned to the onset of running, predictive licking, or reward delivery were obtained for each neuron. Null distributions were created by shuffling firing rate curves and averaging across n = 1000 shuffles. A neuron was defined as having a significant tuning relative to a particular behavioral event if its firing rate exceeded the 99th percentile of the null distribution for at least 0.1s within −1 to 1 s window around the event.

##### Interneuron clustering

Interneurons were identified based on their spike waveforms and their auto-correlograms. Principle component analysis (PCA) was performed on the matrix formed by the normalized theta phase histograms of all the interneurons. The number of PCA components was selected such that the accumulated explained variance reached 95% unless the number of components was larger than 30. The same treatment was done on the matrix formed by the auto-correlograms of all the interneurons. The PCA components of the theta phase histograms, the auto-correlograms, and the estimation of the depth of interneurons relative to the center of the pyramidal layer were features used to perform K-means clustering of interneurons. The number of clusters was determined by the evalclusters function in Matlab, and then went through visual inspection. The putative PV basket cell cluster was identified based on similarities with well-established firing phenotypes of this cell group^15,18^. The estimation of depths of interneurons and pyramidal neurons relative to the center of the pyramidal layer was calculated following the method in Mizuseki et.al., 2011^42^.

##### Model Overview

We developed a computational model to simulate the population dynamics of the hippocampal CA1 region, focusing specifically on the interplay between pyramidal cells with interneurons and nonlinear integration of neuronal inputs. The primary goal of this model is to replicate the observed up- and down-cell dynamics within the pyramidal cell population and to explore the modulatory effects of parvalbumin (PV) and somatostatin (SST) interneurons on functionally defined subpopulations of pyramidal neurons.

The model was developed and implemented using Python version 3.11, with core numerical operations facilitated by NumPy version 1.24. All simulations were run on a Dell Optiplex 3060 workstation with an Intel Core i7-8700 CPU @ 3.20GHz with 32.0 Gb RAM with Windows 11 Enterprise installed. Simulation results were saved as per Neurodata Without Borders (NWB) 2.0 formatting guidelines^43^.

###### Model Structure

Our model is composed of four distinct neuronal populations found within the hippocampal CA1 region^15,28^: CA1 pyramidal cells (n=2000), PV interneurons (n=500), SST interneurons (n=150), and RAD interneurons (n=100). The cell numbers were chosen for a mix of biological realism and computational efficiency^29^. Connections between these populations are established with specific strengths and random probabilities. Each connection is assigned probabilistically (1 or 0), reflecting the likelihood of synaptic connections observed in biological circuits^30^.

Each neuron in the model is described by its firing rate, following the formalism introduced by Wilson and Cowan (1972)^44^. However, unlike traditional rate models that often describe populations, we adapted this framework to model individual neurons within each population. Interneurons (PV, SST, RAD) are modeled as simple point neurons, with their firing rates described by the Wilson and Cowan equations (Eq. 1). Each interneuron’s activity is treated independently to capture the variability observed across the population. The CA1 pyramidal neurons are modeled using a composite function approach, inspired by the compartmental modeling techniques described by Mel and colleagues^31,32^. In this approach, inputs to the distal and apical dendrites are integrated nonlinearly before summing with inputs received at the perisomatic region (Fig. 6a; Eq. 2). This was achieved using a bowl-shaped function (Eq. 3) where one axis was the sum of distal apical inputs, another axis was the sum of proximal apical inputs and the output was in turn the net input of apical currents reaching the soma.

###### External Inputs

The model is driven by external inputs that either increase or decay at the onset of a simulated running event (Eq. 3). We suppose that each neuron within a population receives an equal total number of these inputs. However, the distribution of distal apical run-correlated (n=500), proximal apical run-related (n=500), and reward-related (n=500) inputs for a given cell varies such that a higher number of one type of input corresponds to a lower number of the other type of input. In pyramidal cells, the run-related inputs target the proximal and distal apical dendrites, whereas the reward-related inputs target perisomatic regions (Fig. 6a). The connectivity between these populations were assigned based on defined synaptic strengths and random connectivity probabilities^30^. We model the temporal dynamics of these inputs with an alpha function summed with a baseline firing rate, and the temporal dynamics are depicted in Fig. 6b.

###### Optogenetic Modulation

To mimic optogenetic activation and inactivation experiments of PV and SST interneurons, a positive or negative current for each condition respectively is injected into a fraction of the cells. The parameters for these current injections. To better model these effects and avoid the Gibbs effect, these square pulses exhibit an exponential rise and decay in line with the time constants of light-activated channels^45^.

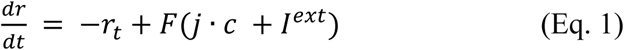

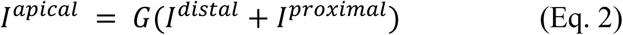

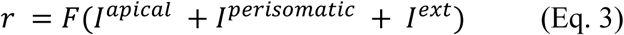

#### Statistics

No statistical methods were used to predetermine sample sizes, but our sample sizes are similar to those reported in previous publications. Data collection was not performed blindly to the conditions of the experiments; however, the data collection and data analysis were performed by two experimentalists to ensure the reproducibility of the results. For optogenetic experiments, stimulation types were randomly determined for each session. We used parametric and non-parametric statistical tests to estimate significant differences between groups: Wilcoxon rank-sum test and two-sample t-test. Both tests are two-sided. All shuffling was done over 1,000 iterations.

## REFERENCES

1. Nielson, D.M., Smith, T.A., Sreekumar, V., Dennis, S., and Sederberg, P.B. (2015). Human hippocampus represents space and time during retrieval of real-world memories. Proc. Natl. Acad. Sci. 112, 11078–11083. 10.1073/pnas.1507104112.

2. Eichenbaum, H. (2017). Time (and space) in the hippocampus. Curr. Opin. Behav. Sci. 17, 65–70. 10.1016/j.cobeha.2017.06.010.

3. Buzsáki, G., and Tingley, D. (2018). Space and Time: The Hippocampus as a Sequence Generator. Trends Cogn. Sci. 22, 853–869. 10.1016/j.tics.2018.07.006.

4. McNaughton, B.L., Battaglia, F.P., Jensen, O., Moser, E.I., and Moser, M.-B. (2006). Path integration and the neural basis of the “cognitive map.” Nat. Rev. Neurosci. 7, 663–678. 10.1038/nrn1932.

5. O’Keefe, J., and Dostrovsky, J. (1971). The hippocampus as a spatial map. Preliminary evidence from unit activity in the freely-moving rat. Brain Res. 34, 171–175. 10.1016/0006-8993(71)90358-1.

6. Robinson, N.T.M., Descamps, L.A.L., Russell, L.E., Buchholz, M.O., Bicknell, B.A., Antonov, G.K., Lau, J.Y.N., Nutbrown, R., Schmidt-Hieber, C., and Häusser, M. (2020). Targeted Activation of Hippocampal Place Cells Drives Memory-Guided Spatial Behavior. Cell 183, 1586–1599.e10. 10.1016/j.cell.2020.09.061.

7. Pastalkova, E., Itskov, V., Amarasingham, A., and Buzsáki, G. (2008). Internally Generated Cell Assembly Sequences in the Rat Hippocampus. Science 321, 1322–1327. 10.1126/science.1159775.

8. Wang, Y., Romani, S., Lustig, B., Leonardo, A., and Pastalkova, E. (2015). Theta sequences are essential for internally generated hippocampal firing fields. Nat. Neurosci. 18, 282–288. 10.1038/nn.3904.

9. Kraus, B.J., Robinson, R.J., White, J.A., Eichenbaum, H., and Hasselmo, M.E. (2013). Hippocampal “Time Cells”: Time versus Path Integration. Neuron 78, 1090–1101. 10.1016/j.neuron.2013.04.015.

10. MacDonald, C.J., Carrow, S., Place, R., and Eichenbaum, H. (2013). Distinct Hippocampal Time Cell Sequences Represent Odor Memories in Immobilized Rats. J. Neurosci. 33, 14607–14616. 10.1523/JNEUROSCI.1537-13.2013.

11. MacDonald, C.J., Lepage, K.Q., Eden, U.T., and Eichenbaum, H. (2011). Hippocampal “Time Cells” Bridge the Gap in Memory for Discontiguous Events. Neuron 71, 737–749. 10.1016/j.neuron.2011.07.012.

12. Taxidis, J., Pnevmatikakis, E.A., Dorian, C.C., Mylavarapu, A.L., Arora, J.S., Samadian, K.D., Hoffberg, E.A., and Golshani, P. (2020). Differential Emergence and Stability of Sensory and Temporal Representations in Context-Specific Hippocampal Sequences. Neuron 108, 984–998.e9. 10.1016/j.neuron.2020.08.028.

13. Eichenbaum, H. (2014). Time cells in the hippocampus: a new dimension for mapping memories. Nat. Rev. Neurosci. 15, 732–744. 10.1038/nrn3827.

14. Heldman, R., Pang, D., Zhao, X., Mensh, B., and Wang, Y. (2023). Time or distance encoding by hippocampal neurons with heterogenous ramping rates. Preprint at Neuroscience, 10.1101/2023.03.12.532295 10.1101/2023.03.12.532295.

15. Klausberger, T., and Somogyi, P. (2008). Neuronal Diversity and Temporal Dynamics: The Unity of Hippocampal Circuit Operations. Science 321, 53–57. 10.1126/science.1149381.

16. Sik, A., Penttonen, M., Ylinen, A., and Buzsaki, G. (1995). Hippocampal CA1 interneurons: an in vivo intracellular labeling study. J. Neurosci. 15, 6651–6665. 10.1523/JNEUROSCI.15-10-06651.1995.

17. Fuchs, E.C., Zivkovic, A.R., Cunningham, M.O., Middleton, S., LeBeau, F.E.N., Bannerman, D.M., Rozov, A., Whittington, M.A., Traub, R.D., Rawlins, J.N.P., et al. (2007). Recruitment of Parvalbumin-Positive Interneurons Determines Hippocampal Function and Associated Behavior. Neuron 53, 591–604. 10.1016/j.neuron.2007.01.031.

18. Royer, S., Zemelman, B.V., Losonczy, A., Kim, J., Chance, F., Magee, J.C., and Buzsáki, G. (2012). Control of timing, rate and bursts of hippocampal place cells by dendritic and somatic inhibition. Nat. Neurosci. 15, 769–775. 10.1038/nn.3077.

19. Stark, E., Eichler, R., Roux, L., Fujisawa, S., Rotstein, H.G., and Buzsáki, G. (2013). Inhibition-Induced Theta Resonance in Cortical Circuits. Neuron 80, 1263–1276. 10.1016/j.neuron.2013.09.033.

20. Korotkova, T., Fuchs, E.C., Ponomarenko, A., von Engelhardt, J., and Monyer, H. (2010). NMDA Receptor Ablation on Parvalbumin-Positive Interneurons Impairs Hippocampal Synchrony, Spatial Representations, and Working Memory. Neuron 68, 557–569. 10.1016/j.neuron.2010.09.017.

21. Leão, R.N., Mikulovic, S., Leão, K.E., Munguba, H., Gezelius, H., Enjin, A., Patra, K., Eriksson, A., Loew, L.M., Tort, A.B.L., et al. (2012). OLM interneurons differentially modulate CA3 and entorhinal inputs to hippocampal CA1 neurons. Nat. Neurosci. 15, 1524– 1530. 10.1038/nn.3235.

22. Hurtado-Zavala, J.I., Ramachandran, B., Ahmed, S., Halder, R., Bolleyer, C., Awasthi, A., Stahlberg, M.A., Wagener, R.J., Anderson, K., Drenan, R.M., et al. (2017). TRPV1 regulates excitatory innervation of OLM neurons in the hippocampus. Nat. Commun. 8, 15878. 10.1038/ncomms15878.

23. Müller, C., and Remy, S. (2014). Dendritic inhibition mediated by O-LM and bistratified interneurons in the hippocampus. Front. Synaptic Neurosci. 6. 10.3389/fnsyn.2014.00023.

24. Lovett-Barron, M., Turi, G.F., Kaifosh, P., Lee, P.H., Bolze, F., Sun, X.-H., Nicoud, J.-F., Zemelman, B.V., Sternson, S.M., and Losonczy, A. (2012). Regulation of neuronal input transformations by tunable dendritic inhibition. Nat. Neurosci. 15, 423–430. 10.1038/nn.3024.

25. Li, N., Chen, S., Guo, Z.V., Chen, H., Huo, Y., Inagaki, H.K., Chen, G., Davis, C., Hansel, D., Guo, C., et al. (2019). Spatiotemporal constraints on optogenetic inactivation in cortical circuits. eLife 8, e48622. 10.7554/eLife.48622.

26. Mahn, M., Gibor, L., Patil, P., Cohen-Kashi Malina, K., Oring, S., Printz, Y., Levy, R., Lampl, I., and Yizhar, O. (2018). High-efficiency optogenetic silencing with soma-targeted anion-conducting channelrhodopsins. Nat. Commun. 9, 4125. 10.1038/s41467-018-06511-8.

27. Diba, K., Amarasingham, A., Mizuseki, K., and Buzsáki, G. (2014). Millisecond Timescale Synchrony among Hippocampal Neurons. J. Neurosci. 34, 14984–14994. 10.1523/JNEUROSCI.1091-14.2014.

28. Topolnik, L., and Tamboli, S. (2022). The role of inhibitory circuits in hippocampal memory processing. Nat. Rev. Neurosci. 23, 476–492. 10.1038/s41583-022-00599-0.

29. Bezaire, M.J., and Soltesz, I. (2013). Quantitative assessment of CA1 local circuits: Knowledge base for interneuron-pyramidal cell connectivity: Quantitative Assessment Of Ca1 Local Circuits. Hippocampus 23, 751–785. 10.1002/hipo.22141.

30. Booker, S.A., and Vida, I. (2018). Morphological diversity and connectivity of hippocampal interneurons. Cell Tissue Res. 373, 619–641. 10.1007/s00441-018-2882-2.

31. Poirazi, P., and Mel, B.W. (2001). Impact of Active Dendrites and Structural Plasticity on the Memory Capacity of Neural Tissue. Neuron 29, 779–796. 10.1016/S0896-6273(01)00252-5.

32. Behabadi, B.F., and Mel, B.W. (2014). Mechanisms underlying subunit independence in pyramidal neuron dendrites. Proc. Natl. Acad. Sci. 111, 498–503. 10.1073/pnas.1217645111.

33. Lee, S.-H., Marchionni, I., Bezaire, M., Varga, C., Danielson, N., Lovett-Barron, M., Losonczy, A., and Soltesz, I. (2014). Parvalbumin-Positive Basket Cells Differentiate among Hippocampal Pyramidal Cells. Neuron 82, 1129–1144. 10.1016/j.neuron.2014.03.034.

34. Valero, M., Cid, E., Averkin, R.G., Aguilar, J., Sanchez-Aguilera, A., Viney, T.J., Gomez-Dominguez, D., Bellistri, E., and De La Prida, L.M. (2015). Determinants of different deep and superficial CA1 pyramidal cell dynamics during sharp-wave ripples. Nat. Neurosci. 18, 1281–1290. 10.1038/nn.4074.

35. Taniguchi, H., He, M., Wu, P., Kim, S., Paik, R., Sugino, K., Kvitsani, D., Fu, Y., Lu, J., Lin, Y., et al. (2011). A Resource of Cre Driver Lines for Genetic Targeting of GABAergic Neurons in Cerebral Cortex. Neuron 71, 995–1013. 10.1016/j.neuron.2011.07.026.

36. Madisen, L., Zwingman, T.A., Sunkin, S.M., Oh, S.W., Zariwala, H.A., Gu, H., Ng, L.L., Palmiter, R.D., Hawrylycz, M.J., Jones, A.R., et al. (2010). A robust and high-throughput Cre reporting and characterization system for the whole mouse brain. Nat. Neurosci. 13, 133–140. 10.1038/nn.2467.

37. Berényi, A., Somogyvári, Z., Nagy, A.J., Roux, L., Long, J.D., Fujisawa, S., Stark, E., Leonardo, A., Harris, T.D., and Buzsáki, G. (2014). Large-scale, high-density (up to 512 channels) recording of local circuits in behaving animals. J. Neurophysiol. 111, 1132–1149. 10.1152/jn.00785.2013.

38. Hazan, L., Zugaro, M., and Buzsáki, G. (2006). Klusters, NeuroScope, NDManager: A free software suite for neurophysiological data processing and visualization. J. Neurosci. Methods 155, 207–216. 10.1016/j.jneumeth.2006.01.017.

39. Pachitariu, M., Steinmetz, N., Kadir, S., Carandini, M., and Kenneth D., H. (2016). Kilosort: realtime spike-sorting for extracellular electrophysiology with hundreds of channels (Neuroscience) 10.1101/061481.

40. Markus, E.J., Barnes, C.A., McNaughton, B.L., Gladden, V.L., and Skaggs, W.E. (1994). Spatial information content and reliability of hippocampal CA1 neurons: Effects of visual input. Hippocampus 4, 410–421. 10.1002/hipo.450040404.

41. Belluscio, M.A., Mizuseki, K., Schmidt, R., Kempter, R., and Buzsaki, G. (2012). Cross-Frequency Phase-Phase Coupling between Theta and Gamma Oscillations in the Hippocampus. J. Neurosci. 32, 423–435. 10.1523/JNEUROSCI.4122-11.2012.

42. Mizuseki, K., Diba, K., Pastalkova, E., and Buzsáki, G. (2011). Hippocampal CA1 pyramidal cells form functionally distinct sublayers. Nat. Neurosci. 14, 1174–1181. 10.1038/nn.2894.

43. Rübel, O., Tritt, A., Ly, R., Dichter, B.K., Ghosh, S., Niu, L., Baker, P., Soltesz, I., Ng, L., Svoboda, K., et al. (2022). The Neurodata Without Borders ecosystem for neurophysiological data science. eLife 11, e78362. 10.7554/eLife.78362.

44. Wilson, H.R., and Cowan, J.D. (1972). Excitatory and Inhibitory Interactions in Localized Populations of Model Neurons. Biophys. J. 12, 1–24. 10.1016/S0006-3495(72)86068-5.

45. Nagel, G., Szellas, T., Huhn, W., Kateriya, S., Adeishvili, N., Berthold, P., Ollig, D., Hegemann, P., and Bamberg, E. (2003). Channelrhodopsin-2, a directly light-gated cation-selective membrane channel. Proc. Natl. Acad. Sci. 100, 13940–13945. 10.1073/pnas.1936192100.

